# Lung tumor-infiltrating T_reg_ have divergent transcriptional profiles and function linked to checkpoint blockade response

**DOI:** 10.1101/2022.12.13.520329

**Authors:** Arbor G. Dykema, Jiajia Zhang, Boyang Zhang, Laurene S. Cheung, Zhen Zeng, Christopher M. Cherry, Taibo Li, Justina X. Caushi, Marni Nishimoto, Sydney Connor, Zhicheng Ji, Andrew J. Munoz, Wenpin Hou, Wentao Zhan, Dipika Singh, Rufiaat Rashid, Marisa Mitchell-Flack, Sadhana Bom, Ada Tam, Nick Ionta, Yi Wang, Camille A. Sawosik, Lauren E. Tirado, Luke M. Tomasovic, Derek VanDyke, Jamie B. Spangler, Valsamo Anagnostou, Stephen Yang, Jonathan Spicer, Roni Rayes, Janis Taube, Julie R. Brahmer, Patrick M. Forde, Srinivasan Yegnasubramanian, Hongkai Ji, Drew M. Pardoll, Kellie N. Smith

**Affiliations:** Bloomberg∼Kimmel Institute for Cancer Immunotherapy, Baltimore, MD, USA; The Mark Foundation Center for Advanced Genomics and Imaging, Baltimore, MD, USA; Sidney Kimmel Comprehensive Cancer Center, Baltimore, MD, USA; Department of Biostatistics, Johns Hopkins Bloomberg School of Public Health; Baltimore, MD, USA; CM Cherry Consulting, Baltimore, MD, USA; Department of Biostatistics and Bioinformatics, Duke University School of Medicine, Durham, NC, 27710, USA; Department of Chemical and Biomolecular Engineering, Baltimore, MD, USA; Translational Tissue Engineering Center, Baltimore, MD, USA; Department of Biomedical Engineering, Baltimore, MD, USA; Department of Surgery, McGill University, Montreal, Canada

## Abstract

Regulatory T cells (T_reg_) are conventionally viewed to suppress endogenous and therapyinduced anti-tumor immunity; however, their role in modulating responses to immune checkpoint blockade (ICB) is unclear. In this study, we integrated single-cell RNAseq/TCRseq of >73,000 tumor-infiltrating T_reg_ (TIL-T_reg_) from anti-PD-1-treated and treatment naive non-small cell lung cancers (NSCLC) with single cell analysis of tumor-associated antigen (TAA)-specific T_reg_ derived from a murine tumor model. We identified 10 subsets of human TIL-T_reg_, most of which have high concordance with murine TIL-T_reg_ subsets. Notably, one subset selectively expresses high levels of OX40 and GITR, whose engangement by cognate ligand mediated proliferative programs and NF-kB activation, as well as multiple genes involved in T_reg_ suppression, in particular LAG3. Functionally, the OX40^hi^GITR^hi^ subset in the most highly suppressive *ex vivo* and T_reg_ expression of OX40, GITR and LAG3, correlated with resistance to PD-1 blockade. Surprisingly, in the murine tumor model, we found that virtually all TIL-T_reg_ expressing T cell receptors that are specific for TAA fully develop a distinct Th1-like signature over a two-week period after entry into the tumor, down-regulating FoxP3 and up-regulating expression of *TBX21 (*Tbet), IFNγ and certain pro-inflammatory granzymes. Application of a gene score from the murine TAA-specific Th1-like T_reg_ subset to the human single-cell dataset revealed a highly analogous subcluster that was enriched in anti-PD-1 responding tumors. These findings demonstrate that TIL-T_reg_ partition into multiple distinct transcriptionally-defined subsets with potentially opposing effects on ICB-induced anti-tumor immunity and suggest that TAA-specific TIL-T_reg_ may positively contribute to anti-tumor responses.

**One-Sentence Summary:** We define 10 subsets of lung cancer-infiltrating regulatory T cells, one of which is highly suppressive and enriched in anti-PD-1 non-responders and the other is Th1-like and is enriched in PD-1 responders.

## Main Text

Immune checkpoint blockade (ICB) induces durable clinical responses in a subset of patients with non-small cell lung cancer (NSCLC), however most tumors do not respond (*1*). ICB efficacy relies on endogenous tumor-reactive T cells for tumor eradication. In our ongoing efforts to understand mechanisms of ICB response vs resistance, we recently reported on the transcriptional programming of CD8+ tumor-infiltrating lymphocytes (TIL) in resectable lung cancers treated with neoadjuvant ICB (*2*). We found that the breadth and frequency of CD8+ TIL recognizing antigens derived from tumor somatic mutations (neoantigens) was not associated with pathologic response, thereby suggesting the existence of immunosuppressive mechanisms that can thwart otherwise effective anti-tumor immunity.

One candidate immune inhibitory cell population in the tumor microenvironment (TME) is the regulatory T cell (T_reg_), characterized by expression of CD4 and the forkhead box protein P3 (FoxP3). These cells maintain systemic immune homeostasis by regulating peripheral tolerance and mitigating autoimmune disease (*3*) through a variety of immunosuppressive mechanisms (*4*). T_reg_ depletion experiments in mice demonstrate that this population can potently suppress anti-tumor effector T cell responses and prevent tumor clearance by endogenous tumor-specific effector T cells (*5, 6*). T_reg_ representation among total TIL in diverse murine and human cancers is generally much higher than in corresponding normal tissue and blood and increased tumor infiltration by T_reg_ expressing specific genes was recently correlated with poor prognosis in NSCLC (*7, 8*). These findings have led to multiple clinical approaches to selectively deplete or inhibit T_reg_ in tumors, but none to date have been successful, possibly because of lack of T_reg_ specificity or failure to inhibit key functionally relevant T_reg_ subsets. Thus, an understanding of the phenotype and specificity of tumor-infiltrating lymphocyte T_reg_ (TIL-T_reg_) is critical to target them for effective immunotherapy.

Contrary to conventional CD4 T cells (T_conv_), whose self-specific repertoire is negatively selected, T_reg_ are selected for expression of T cell receptors (TCR) that recognize self-antigen, thereby allowing them to maintain self-tolerance via targeted immune suppression. Under certain circumstances of pathologic autoimmunity, T_reg_ can lose FoxP3 and differentiate to acquire a T_conv_ program (so-called ‘ex-T_reg_’) and thereby exacerbate autoimmunity via interferon-_γ_ (IFN_γ_) production and recruitment of Th1 cells (*9*). While historically T_reg_ were dichotomized into thymic (natural T_reg_) vs. peripheral (induced T_reg_) in origin, single-cell RNA-sequencing (scRNA-seq) technology has enabled more refined analysis of these cells to reveal a level of T_reg_ heterogeneity that was not previously appreciated (*10*–*16*). To study the transcriptional programming and function of TIL-T_reg_ at refined resolution, we employed single cell RNA-sequencing of TIL-T_reg_ harvested from resected human lung cancers (and adjacent normal lung) from untreated patients, those treated with anti-PD-1 and, in parallel, integrated these data with an analysis of a murine tumor model wherein tumor-associated antigen (TAA)-specific T_reg_ were tracked. We identified two distinct T_reg_ subsets displaying opposite association with ICB response. One is an activated, highly suppressive T_reg_ subset, characterized by a unique tumor necrosis factor superfamily (TNFRSF) expression pattern and high levels of genes encoding multiple suppressive and tumor-homing molecules, that associates with ICB non-responsiveness. The other is a unique population of TAA-specific T_reg_, defined in murine tumors, that exhibit certain characteristics of ‘ex-T_reg_’, with Foxp3 downregulation and an acquired Th1-like effector program. Using computational gene score homology analysis, we identified an analogous population in human lung cancers that was enriched in ICB responding tumors. These findings reveal a previously unappreciated diverse substructure of TIL-T_reg_ that consists of distinct subsets that can either inhibit or enhance anti-tumor responses, defining potential T_reg_ targets for therapeutic modulation.

## Results

### T_reg_ transcriptional programs in resected lung cancers treated with ICB

To better understand the transcriptional programs of TIL-T_reg_ subsets and their potential role in ICB responsiveness, we analyzed a dataset of coupled single cell TCRseq/RNAseq on CD3+ TIL isolated from resected lung cancers and adjacent normal lung (NL) tissue, derived from patients treated with neoadjuvant PD-1 blockade (NCT02259621) (*17*). We have previously reported on transcriptional programming of neoantigen-specific CD8+ T cells from this dataset (*2*). Of these 15 ICB-treated tumors, six had a major pathologic response (MPR), defined as ≤10% residual tumor at the time of surgery (**Table S1**), which was recently associated with improved overall survival in lung cancers treated with neoadjuvant combination immunotherapy in the CheckMate 816 trial (NCT02998528) (*18*). Per standard convention, we refer to tumors with MPR as responders (R) and tumors without MPR as non-responders (NR). Additionally, coupled single cell TCRseq/RNAseq was performed prospectively on CD3+ T cells from tumor and adjacent NL from 10 treatment-naive patients with resectable lung cancer (**Table S2**; **Fig. S1a**). After stringent quality-control filtering, 851,674 CD3+ T cells passed quality control and were integrated to obtain broad CD4, and CD8 T cell subsets as visualized by Uniform manifold approximation and projection (UMAP) dimensionality reduction (**Fig. S1b**). Refined clustering of 375,001 CD8-, CD4+ T cells identified 11 subclusters, with 3 distinct T_reg_ clusters (**Fig. S1c**). Computational selection of FoxP3^+^CD8^-neg^ cells identified 73,882 T_reg_. This large number of individual transcriptomes – the largest reported from a single integrated analysis - allowed us to cluster the TIL-T_reg_ with even higher resolution, thus identifying 10 distinct T_reg_ subclusters (**Fig. 1a; Table S3**). Four ‘Activated’ clusters were identified based on high expression of *IL2RA, CCR8*, and *ICOS*, which are broadly associated with T_reg_ activation, and expression of published gene sets induced by TCR activation (KEGG) and IL-2 signaling (BIOCARTA) (**Fig. 1b, Fig. S1d**). Cluster-defining genes and select functional markers were visualized (**Fig. 1b, c**). Notably, Activated (1)/ OX40^hi^GITR^hi^ had a transcriptional program consistent with a unique activated state, as evidenced by high selective expression of the TNFSF members *TNFRSF4* (OX40) and *TNFRSF18* (GITR) (**Figs. 1b-c, S1e-f**) (*7*). A number of additional activation genes were shared with one of the other activated clusters, including *LAG-3* (Activated (2); **Fig. S1g**), and *TNFRSF9* (41BB), *ICOS*, and *CCR8* (Activated (3)). In contrast, there was a LN homing cluster (*SELL, S1PR1)*, two ‘resting’ clusters based on expression of *SESN3* and gene sets associated with resting T_reg_ (**Fig. 1b, c, Fig. S1d**), and an ‘inactive’ population with overall low differential gene expression or gene set enrichment analysis (GSEA) association (**Fig. 1b; Fig. S1d**). Other distinct clusters included a type I IFN-responsive cluster enriched for IFN-inducible genes such as *IFI6* and *MX1* and expression of an IFN-responsive gene set, and an unusual subset characterized by expression of genes typically expressed by Th1 cells, such as *CCL4 (*MIP-1β*), CCL5* (RANTES), *IFN*_γ_, and multiple granzyme genes typically found in activated CD8 cytotoxic lymphocytes (**Fig. 1b, c; Table S3**). Cell proportion analysis found enriched representation of the Activated (1)/OX40^hi^GITR^hi^ (p=0.0054), Resting (2) (p=8.8E-10), and Activated (4) (p=0.014) clusters in tumor relative to adjacent NL, whereas the LN-homing cluster was enriched in adjacent NL (p=4.5E-08; **Fig. S2a**). The distinct pathologic responses among different patients in the trial allowed us to ascertain whether specific T_reg_ subsets were associated with ICB response, but globally, there was no difference in TIL-T_reg_ subcluster proportions between treated and untreated tumors, nor between R and NR (**Fig. S2b**). Principal component analysis (PCA) of pseudobulk gene expression separated TIL-T_reg_ from those derived from adjacent NL (**Fig. 1d**), indicating distinct global transcriptional programs in tumor-derived vs. adjacent NL T_reg_, but could not distinguish R from NR (**Fig. 1e**) nor treated from treatment-naive (**Fig. S2c**).

**Figure 1.**
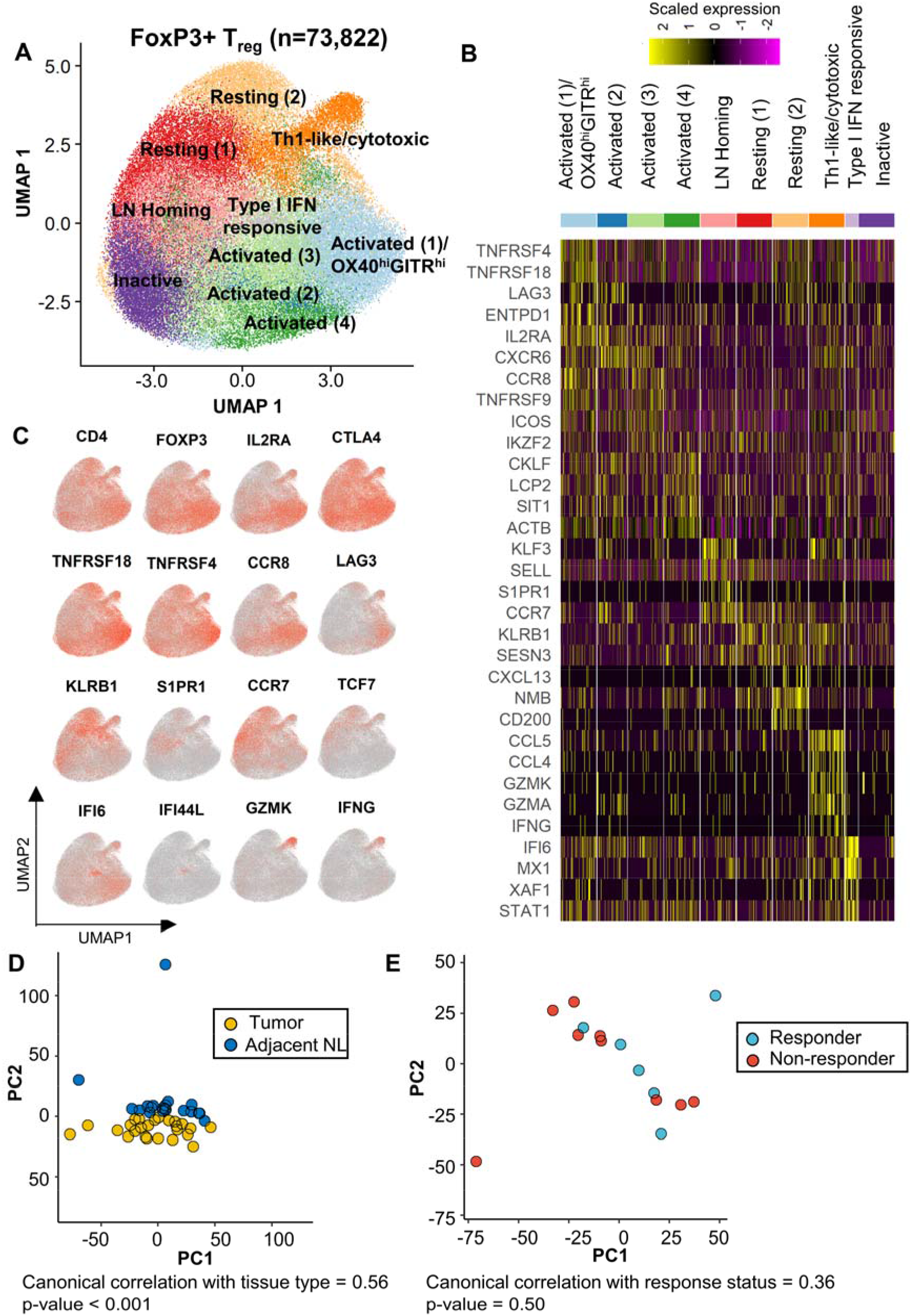
Single cell transcriptomic profiling of T_reg_ in treatment naïve and neoadjuvant anti-PD-1 treated non-small cell lung cancers. Coupled single-cell (sc) RNA-seq/TCR-seq was performed on T cells isolated from resected tumor (n=15), adjacent normal lung (NL; n=12), tumor draining lymph node (TDLN; n=3), and a resected brain metastasis (n=1) from NSCLC patients treated with two doses of neoadjuvant anti-PD-1 as well as resected tumor (n=10) and paired adjacent normal lung (n=8) from treatment-naïve patients with NSCLC. A, 2D UMAP projection of the expression profiles of the 73,882 T_reg_ that passed QC. T_reg_ subsets, defined by 10 unique clusters, are annotated and marked by color code. B, Relative expression of top differential genes for each cluster is visualized on a heatmap. 3,000 cells (or all cells in the cluster if cluster size <3,000 cells) were randomly sampled from each cluster for visualization. C, The expression of canonical T_reg_ subset marker genes and cell subset selective genes were visualized in red-scale using UMAP projection. D, Principal component analysis (PCA) and canonical correlation of pseudobulk gene expression for individual tumor (yellow, n=25) and adjacent NL (dark blue, n=20) samples. E, Principal component analysis (PCA) and canonical correlation of pseudobulk gene expression for individual non-responder (red, n=9) and responder (blue, n=6) tumors.

### OX40^hi^GITR^hi^ T_reg_ are highly suppressive and associated with ICB resistance

While OX40, GITR, and 4-1BB have been targeted with agonist antibodies based on their co-stimulatory roles on T_conv_, they are also expressed by TIL-T_reg_ (*19*), in which their role has yet to be clarified. Among all the TIL-T_reg_, the Activated (1)/OX40^hi^GITR^hi^ cluster exhibited markedly increased expression of multiple genes encoding molecules that mediate suppressive activity, such as *FoxP3, IL2RA, LAG3, ENTPD1 (*CD39*), EBI3*, and *LAYN* (*7, 20*–*23*) when compared to all other T_reg_ clusters, suggesting a highly immunosuppressive phenotype (**Fig. 1b; Table S4)**. To determine whether the Activated (1)/OX40^hi^GITR^hi^ T_reg_ displayed functionally superior suppressive capacity, we sorted OX40^hi^GITR^hi^ and OX40^low^GITR^low^ T_reg_ from ICB-treated tumors to test their ability to suppress T_conv_ cells *ex vivo*. Indeed, OX40^hi^GITR^hi^ T_reg_ were markedly more suppressive than OX40^low^GITR^low^ T_reg_ (**Fig. 2a; Fig S3**). While the proportion of individual T_reg_ subclusters was not statistically associated with ICB response (**Fig. S2b**), we hypothesized that T_reg_ activation status could be, owing to the enrichment of the Activated (1)/OX40^hi^GITR^hi^ cluster in TIL compared to adjacent NL T_reg_ (**Figs. S2a, S3a**). We therefore developed an “activated” T_reg_ score, derived from the top differentially expressed genes in Activated (1)/OX40^hi^GITR^hi^: *LAG3, TNFRSF4* (OX40), and *TNFRSF18* (GITR) (**Figs. 2b, S1e-g, Table S4**). TIL-T_reg_ expressing the activated T_reg_ score were localized within the Activated (1)/OX40^hi^GITR^hi^ cluster **(Figs. 2c, S4b)** and significantly more enriched in NR vs R tumors (p=0.038; **Figs. 2c,d)**. Interestingly, treatment-naïve tumors had high representation of T_reg_ with a high activation score, similar to ICB NRs (p=0.670), suggesting that contraction of this subset was a consequence of successful ICB treatment in responding tumors. Of note and in contrast, the unusual Th1-like/cytotoxic cluster (**Figs. 1a,b**) was not enriched in NR tumors. There was no difference in frequency of T_reg_ with a high activation score in adjacent NL among the different groups (p=0.620), indicating that the difference between NR and R is specific to the tumor microenvironment (**Fig. 2d**). Enrichment of “activated” T_reg_ in NR tumors could be either due to differentiation into a more activated, suppressive state, better maintenance of that state, and/or prevention of differentiation out of that state. To address this question, we assessed TCR clonal sharing between each subcluster, based on the notion that clones shared between and among clusters can serve as lineage tracking markers and thus reflect developmental connectivity. Indeed, of 1,047 clones found in more than one cluster, 699 (66.8%) were shared between Activated (1)/OX40^hi^GITR^hi^ and Activated (3) (**Fig. 2e**). Next, we performed diffusion trajectory and RNA velocity analysis on T_reg_ clones shared in the Activated (1)/OX40^hi^GITR^hi^ and Activated (3) clusters to define the developmental relationships between these clusters in NR vs. R tumors. This analysis showed that TIL-Treg within R tumors differentiate out of Activated (1)/OX40^hi^/GITR^hi^ into Activated (3), contrasting with what is seen in NR tumors, in which the TIL-Treg developmental trajectory is into and within the Activated (1)/OX40^hi^GITR^hi^ cluster with essentially no differentiation out of that state (**Fig. 2f**). Taken together, these findings indicate the paucity of Activated (1)/OX40^hi^GITR^hi^ cells in R tumors is due to a net differentiation flux out of this subset, in contrast to NR (**Fig. 2f)** (*24*). In support of this, the proportion of Activated (1) relative to Activated (3) was significantly decreased in R tumors relative to NR (p=0.0115) and treatment-naïve (p=0.0136; **Fig. 2g**)

**Figure 2.**
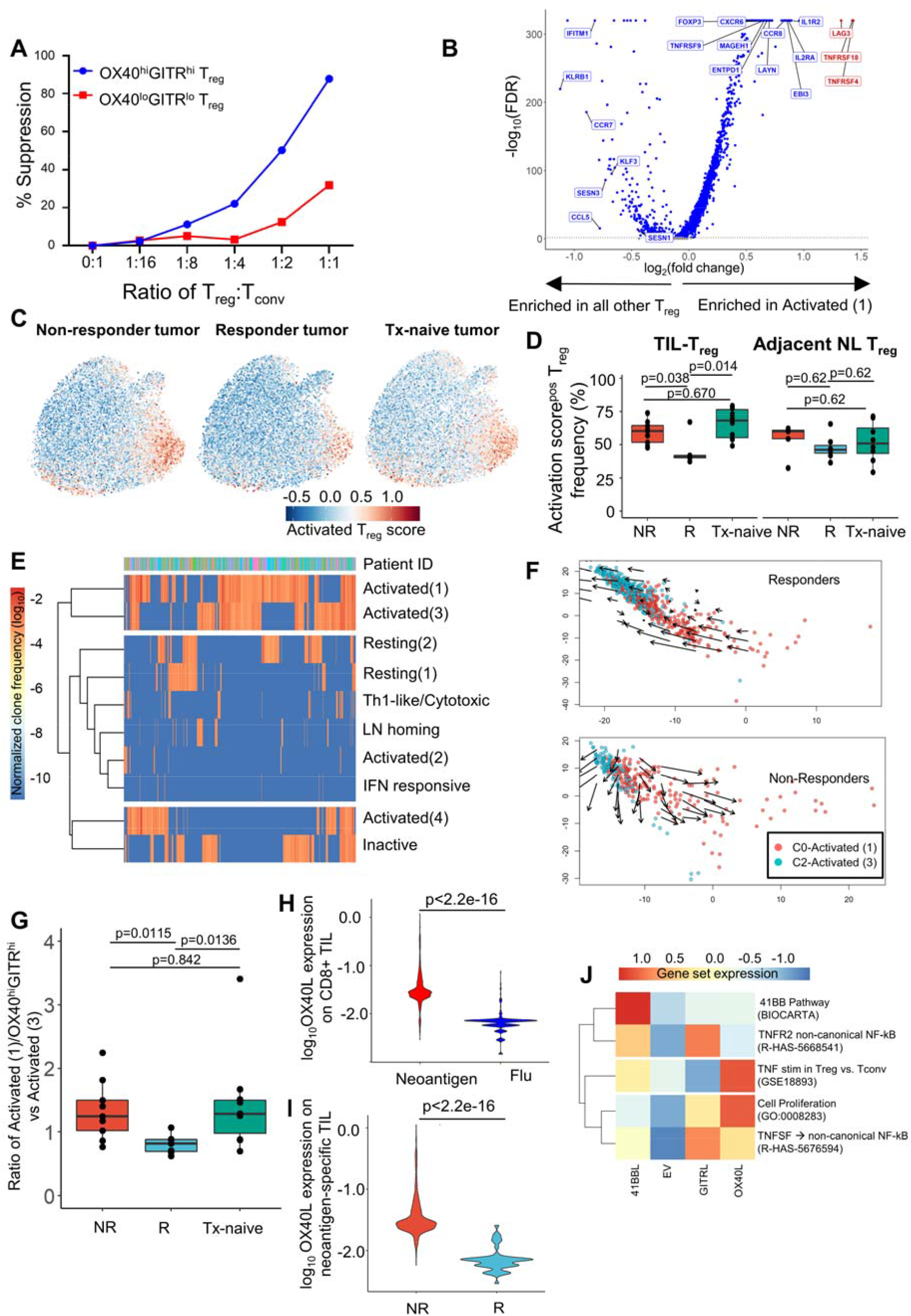
Activated, OX40^hi^GITR^hi^ T_reg_ are functionally suppressive and associate with non-response to PD-1 blockade. A, Functional analysis of OX40^lo^GITR^lo^ and OX40^hi^GITR^hi^ TIL-T_reg_-mediated suppression of conventional CD4^+^T cell (T_conv_) proliferation. % suppression was calculated by: ((% divided of T_conv_ cells alone - % divided of T_conv_ cells cultured with T_reg_)/ % divided of T_conv_ cells alone)*100. B, Volcano plot showing differential expression between the Activated (1)/OX40^hi^GITR^hi^ cluster (right) vs. all other T_reg_ (left). Each dot represents one gene. FDR<0.05 is considered significant (blue/red dots). LAG3, TNFRSF18 and TNFRSF4 represent the top 3 most differentially expressed genes in the Activated (1)/OX40^hi^GITR^hi^ T_reg_ (red dots). C, Overlay of the Activated T_reg_ score on the TIL-T_reg_ UMAP for each response/treatment group. Red indicates higher expression; blue indicates lower expression. D, The frequency of T_reg_ with a high T_reg_ activation score is shown for tumor (left) and adjacent NL (right) from non-responders (NR; red), responders (R; blue), and treatment-naive patients (Tx-naive; green). Comparisons were performed at the individual patient level using Wilcoxon rank test. E, Clonotype sharing pattern across T_reg_ subsets. The frequency of T_reg_ TCR clones that were detected in at least two T_reg_ clusters were calculated and shown on a heatmap. F, Diffusion plot with RNA velocity for the Activated (1)/OX40^hi^GITR^hi^ and Activated (3) clusters (among which most clonotype sharing was observed). G, Boxplots showing the relative proportion of Activated (1)/OX40^hi^GITR^hi^ and Activated (3) clusters by responders (blue), non-responders (red), and treatment-naïve status (green). H, A violin plot shows TNFSF4 (OX40L) expression by neoantigen-specific (red) and flu-specific (blue) CD8 TIL from the same neoadjuvant-t5reated patients. I, A violin plot compares the TNFSF4 (OX40L) expression by neoantigen-specific CD8+ TIL between non-responding (red)and responding (blue) tumors. J, Gene set enrichment analysis to evaluate differing biological functions of ex vivo OX40L, 41BBL, and GITRL agonism in sorted human TIL-T_reg_.

Beyond being markers for a highly suppressive T_reg_, OX40, GITR, and 4-1BB - the three TNFRSF members most highly expressed by the Activated (1)/ OX40^hi^/GITR^hi^ subset, likely play active roles in their function (*24*) that would be dependent on expression of their cognate ligands within the TME. We therefore broadly queried the expression of TNFR-ligands on all intratumoral T cell and myeloid populations to define specific cell types that may be interacting with the OX40 and GITR-expressing T_reg_. We found that in our treated and untreated NSCLC, GITR-ligand (GITRL, *TNFSF18*) was very lowly expressed across all cell types in the TME: 4-1BB-ligand (4-1BBL, *TNFSF9*) was broadly expressed on both myeloid and T cells: and OX40-ligand (OX40L, *TNFSF4*) was largely expressed on a limited number of T cells, most notably on CD8+ T cells (**Fig. S4c)**. Given that expression of OX40L was restricted to CD8 T cells, we queried our previously-published flu- and neoantigen-specific CD8+ TIL in this patient cohort (*2*). There was significantly higher expression of OX40L on neoantigen-vs. flu-TIL (**Fig. 2h**, p< 2.2E-16). Even more remarkably, the expression by neoantigen-specific TIL was much higher on neoantigen-specific CD8+ TIL from NR compared to R tumors (**Fig. 2i**, p<2.2E-16). There was no significant difference in expression in 41BB-ligand on neoantigen-vs. flu-TIL, regardless of response status (**Fig. S4d)**.

To assess the effect of TNFR signaling on T_reg_ transcriptional programs, we generated HEK-293T cell lines expressing each of the three TNFR ligands (OX40L, GITRL, and 41BBL; **Fig. S5a**). Sorted T_reg_ from two NR tumors were co-cultured with the ligand-expressing 293T cells, followed by single cell TCRseq/RNAseq of the T_reg_. In each ligand-stimulated condition, the non-canonical NFkB pathway was enriched (R-HAS-5676594, GSE18893; **Fig. 2j, Table S5-7**). Specifically, 4-1BB agonism initiated its unique gene program (BIOCARTA_41BB_PATHWAY; **Table S5**): 4-1BB and GITR agonism showed enrichment for a gene set associated with TNFR2 signaling (R-HAS-5668541; **Table S5, S6**) (*25*); and OX40 agonism strongly upregulated a cell proliferation gene set (GO:0008283; **Table S7**; **Fig. S2j**). Given the particularly high expression of OX40L on neoantigen-specific CD8+ T cells in NR patients (**Fig. 2i**), our findings suggest that they may be providing a signal to expand this suppressive OX40^hi^GITR^hi^ T_reg_ subset, thereby subverting their anti-tumor activity.

### Tumor antigen-specific T_reg_ acquire a Th1-like transcriptional signature in an ICB responsive murine tumor model

In direct contrast to T_conv_ cells, T_reg_ are selected to recognize self-antigen (*26*). Indeed, naturally occurring T_reg_ (nT_reg_) are positively selected in the thymus by a broad set of self-antigens expressed in thymic epithelial cells (*27*–*29*). In order to model the biology of tumor-associated self-antigen (TAA)-specific TIL-T_reg_, we developed a mouse adoptive transfer model of T_reg_ with fixed specificity. We crossed the B6.Cg-Tg(TcrLCMV) TCR transgenic line that has a CD4+ TCR (termed SMARTA) specific for the H2-I-A^b^-restricted LCMV GP_61-80_ epitope (*30*), to mice that have expression of the LCMV glycoprotein (GP) controlled by the rat insulin promoter (RIP-GP) (*31, 32*). It has been well documented that RIP is also active in murine thymic epithelial cells. Consistent with the developmental biology of T_reg,_ virtually no FoxP3+ T_reg_ bearing the SMARTA TCR develop in the absence of GP as a self-antigen, so this mating results in offspring with a much higher frequency of LCMV GP_61-80_ – restricted CD4+FoxP3+ T_reg_ due to thymic expression of GP (*33, 34*). We then crossed these mice to CD45.1;RFP-FoxP3 mice to generate a T_reg_ reporter mouse line with a transgenic TCR of known reactivity. CD45.1-marked transgenic CD25^hi^RFP-FoxP3^+^ T_reg_ expressing the SMARTA TCR alpha and beta chains (Vα2.3-JαDK1 and Vβ8.3-Jβ2.5) were sorted and 2×10^5^ SMARTA TCR^pos^ T_reg_ per mouse were adoptively transferred to recipient B6 CD45.2 mice (**Fig. S6a**). Recipient mice were inoculated with either the MC38 parental line (MC38WT), which is an ICB-responsive tumor, or a MC38-GP-expressing line (*35*). After 14 days of tumor growth, we harvested the tumors and sorted TIL-T_reg_ based on high expression of CD25. The sorted population consisted of a mix of distinguishable tumor-reactive (TR)-T_reg_ that were adoptively transferred (SMARTA TCR^pos^/RFP^pos^) and endogenous T_reg_ (SMARTA TCR^neg^/RFP^neg^) (**Fig. S6b)**. Of note, fewer than 1% of total T_reg_ in the spleen were from the donor SMARTA TCR^pos^ T_reg_ 14 days after adoptive transfer, a proportion similar to previous protocols using adoptively transferred T_conv_ (**Fig. S6c**)(*36*).

Single cell TCRseq/RNAseq was performed on >30,000 murine TIL-T_reg_, which identified five clusters (**Fig. 3a, b; Table S8**). As expected, TR-T_reg_ accumulated in GP-expressing tumors and tumor-draining lymph node (TDLN) to a significantly greater extent than in MC38WT tumors (p=0.0159 and p=0.0317, respectively), which indicates antigen recognition driving subsequent cell activation and proliferation (**Fig. 3c**).

**Figure 3.**
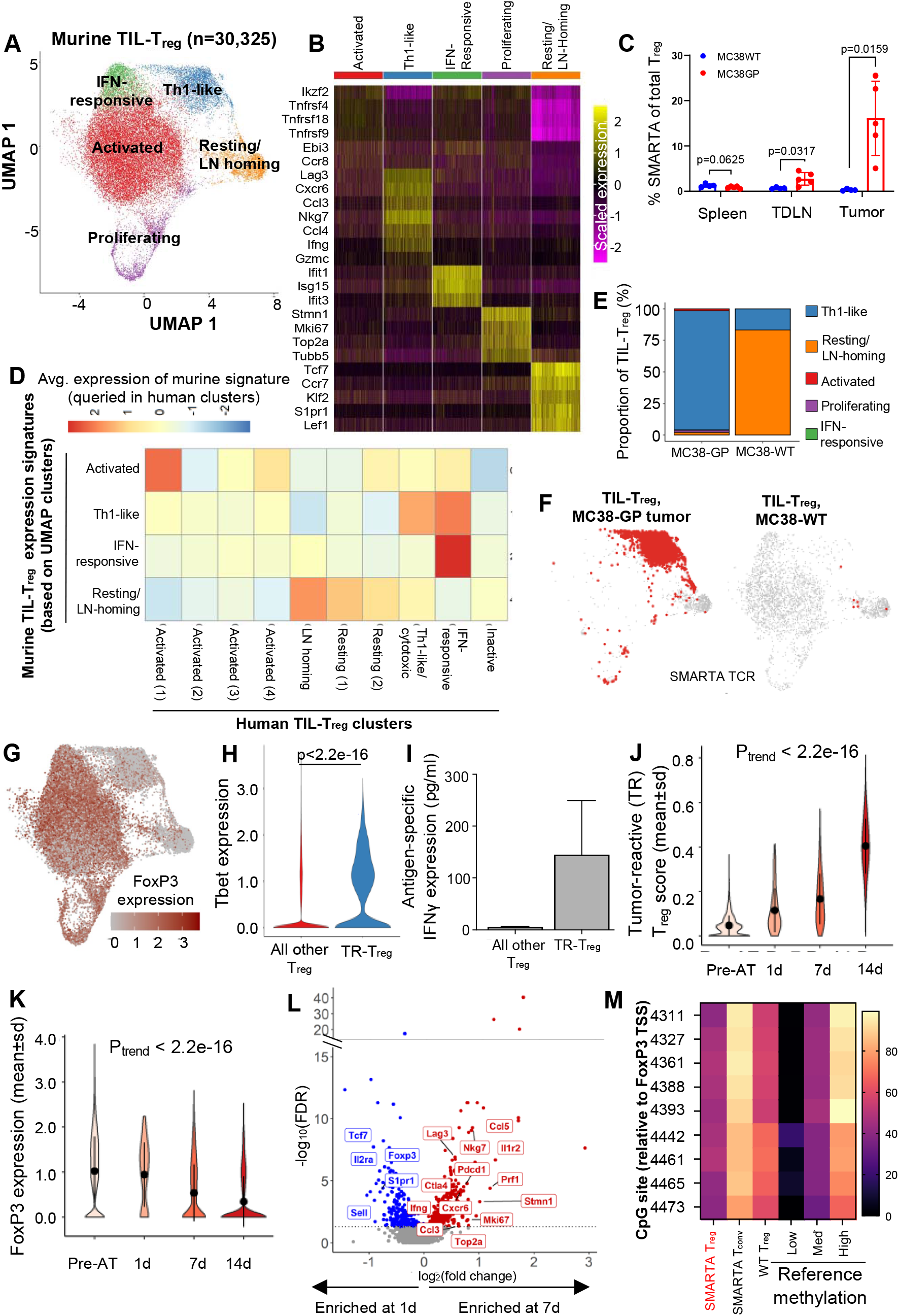
Defining the scRNA-transcriptome associated with T_reg_ tumor antigen-reactivity using a transgenic TCR mouse model. Coupled single-cell (sc) RNA-seq/TCR-seq was performed on T_reg_ isolated from GP-expressing MC38 (MC38-GP) or parental MC38 (MC38WT) tumors, tumor draining inguinal lymph node, and spleens on Day 14 of tumor growth. A, 2D UMAP projection of the expression profiles of the 30,325 T_reg_ that passed QC. T_reg_ subsets, defined by 5 unique clusters, are annotated and marked by color code. B, Relative expression of 3-5 differential genes for each cluster is visualized on a heatmap. C, Comparison of the frequency of SMARTA FoxP3+ T_reg_ of total T_reg_ defined by expression of CD45.1, FoxP3 and the SMARTA TCR between MC38WT (blue) and MC38-GP (red) tumor bearing mice in each tissue compartment. Mean with SEM error bars are shown for each mouse (n=5). P value obtained using paired student’s t-test; ns, non-significant values. D, Homology to human T_reg_ clusters is shown by the average expression of genes scores built on the top 20 differentially expressed genes for each murine cluster queried in human clusters. E, Quantification of cluster designation for all SMARTA clones from WT and GP-expressing MC38. F, Cluster localization for SMARTA clones is shown with SMARTA TCR (red) overlaid on full 2D UMAP (grey) of T_reg_ isolated from MC38-GP (left) or MC38WT (right). G, FoxP3 expression is visualized in red scale using full UMAP projection. H, Tbet expression is shown for TR-T_reg_ (blue) and all other T_reg_ (red) from the GP-expressing MC38 tumor after 14 days in vivo. I, Five thousand TR-T_reg_ and non-TR-T_reg_ were sorted and co-cultured ex vivo with one thousand dendritic cells loaded with the LCMV GP peptide for 48h. Supernatants were harvested and assayed for IFN_γ_ using the Meso Scale Discovery platform. Data are shown as IFN_γ_ production above background, defined as T_reg_ co-cultured with dendritic cells without peptide. J, Violin plot showing the TR-T_reg_ (tumor-reactive T_reg_) score of adoptively-transferred SMARTA TCR^pos^ T_reg_ before adoptive transfer (pre-AT), and at 1, 7, and 14 days of tumor residence. K, Violin plot showing FoxP3 expression by adoptively-transferred SMARTA TCR^pos^ T_reg_ before adoptive transfer (pre-AT), and at 1, 7, and 14 days of tumor residence. P values obtained by Student’s t-test. L, Volcano plots showing differentially expressed genes of SMARTA+ Tregs in day 7 vs day 1. x-axis shows log2(fold change) and y-axis –log10(FDR). Differentially expressed genes higher at day 7 are shown in red, and differentially expressed genes higher in 1 day in blue. M, T_reg_-specific demethylated region (TSDR) methylation profile of sorted SMARTA+CD25^hi^RFP+ T_reg_ from female SMARTA;RIP-GP mice compared to control values (low, med, high). Residue numbers denote individual CpG motifs in references to the transcription initiation site of FoxP3. Bar colors represent 0-100% methylation.

To compare our murine and human TIL-T_reg_ clusters, we used the top 20 genes from each of the murine T_reg_ clusters to develop gene scores. We then queried our human clusters to see which human clusters corresponded most to the murine clusters identified. Remarkably, the murine T_reg_ clusters were highly related to counterpart human TIL-T_reg_ clusters (**Fig. 3d**). Specifically, there was notable homology in the top cluster-defining genes for the resting and LN-homing clusters (*CCR7, TCF7, KLF2, S1PR1*), the IFN-induced cluster (*IFIT3, ISG15, IRF7)*, and the Th1-like cluster (*CCL5, CCL4, IFNG)* (**Fig. 3b, d**; **Table S9)**. Murine TIL-T_reg_ also contained an activated cluster in which the top differentially expressed genes (*TNFRSF4* (OX40), *TNFRSF18* (GITR), *TNFRSF9* (41BB), *IZKF2, EBI3, CCR8*) represent a compilation of several of the human activated clusters and was most analogous to Activated (1)/OX40^hi^GITR^hi^ (**Fig. 3d**). A notable difference was the proliferating cluster. While there are human TIL-T_reg_ with proliferation signatures, they are relatively few and did not segregate out into a separate cluster as they did in the mouse. This is likely because the relative kinetics of human vs murine tumor growth allows for a greater equilibrium to develop over the years of tumor development in humans. We next queried the murine single cell data for co-expression of the SMARTA TCR alpha and beta chain to identify the transcriptional profile of tumor-reactive Treg (TR-T_reg_). Notably, in MC38-GP tumors the TR-T_reg_ did not localize to the Activated (1)/OX40^hi^GITR^hi^ T_reg_ cluster, but rather acquired a transcriptional program characterized by up-regulation of Th1-associated cytokine and pro-inflammatory chemokine genes such as IFNγ, CCL3 (MIP1α), and CCL4 (MIP1β) (**Fig. 3b; Table S10**). In contrast, the rare TR-T_reg_ cells harvested from the MC38WT tumors all resided in the resting/LN-homing cluster (**Fig. 3e, f**), consistent with a lack of antigen stimulation (**Fig. 3b, d**). Because many of the genes associated with the TR-T_reg_ after 14 days’ residence in the tumor are not typically seen in conventional suppressive T_reg_, we assessed the level of FoxP3 expression in each mouse T_reg_ cluster. Indeed, the TR-T_reg_ cluster, while still RFP+ (indicating that the Foxp3 locus had been transcriptionally active) had relatively low FoxP3 mRNA expression compared to the other clusters (**Fig. 3g**). This is consistent with the long half-life of RFP that allows it to be detected after transcription of the linked gene is turned off (*37*). Antigen-dependent loss of FoxP3 expression in a minority of T_reg_ has been observed in models of autoimmunity and is associated with loss of T_reg_ suppressive capacity and acquisition of T_conv_ activity (*9*). To determine if the decreased FoxP3 mRNA was mirrored at the protein level, we sorted RFP+ T_reg_ directly from SMARTA (TR-T_reg_) donor mice and compared intracellular FoxP3 protein expression to TR-T_reg_ isolated from recipient mice after the TR-T_reg_ had been in the MC38-GP expressing tumor for 14 days. Prior to adoptive transfer, an average of 94.2% (range 92.4 – 95.3%) of the sorted population was FoxP3+, while only 22.8% were FoxP3+ after 14 days (**Fig. S6d**). Furthermore, the TR-T_reg_ cluster was highly enriched for Tbet (*TBX21*) expression (**Fig. 3h**), which is a critical transcriptional regulator of the proinflammatory function of Th1 cells as well as Th1-like ex-T_reg_ observed in autoimmune settings (*38*–*42*). Although Th1 cells are known to be proinflammatory, they only kill cells indirectly through activation of nearby phagocytic cells (*43*). In contrast, T_reg_ have been reported to directly kill cells through release of cytotoxic granules containing granzymes and perforin (*44*). Because our murine TR-T_reg_ upregulate some cytotoxic granule molecules and *NKG7* (*45, 46*) (**Fig 3b,h**), we asked whether they could kill MC38-GP tumor cells induced to express MHC2 through TCR:MHC binding. We harvested TIL TR-T_reg_ and endogenous T_reg_ on Day 14 of tumor growth and plated them at varying ratios with IFNγ-induced, MHC2+ MC38-GP tumor cells (**Fig. S7a**). TR-T_reg_ did not exhibit any specific killing against MHC2+ GP-expressing tumor cells (**Fig. S7b**), but they did functionally produce IFNγ in an antigen-specific fashion when co-cultured with LCMV GP_61-80_ peptide-loaded dendritic cells ex vivo (**Fig. 3i**, suggesting they could potentially assist anti-tumor immunity via a helper rather than cytotoxic mechanism. Additionally, the upregulation in the TR-T_reg_ cluster of Tbet, which has a known role in mediating the proinflammatory function of Th1-like T_reg_ (*38*–*42*) (**Fig. 3h**) supports this notion. Taken together, the data demonstrate that tumor antigen-reactive T_reg_ acquire a distinct phenotype consistent with an “ex-T_reg_” Th1-like gene signature, accompanied by loss of *FoxP3* and *CD25* expression and upregulation of *CCL5* and *IFN*_γ_.

To evaluate the kinetics of acquisition of this this Th1-like phenotype, we performed single cell TCRseq/RNAseq on TR-T_reg_ harvested one, seven, and fourteen days post-adoptive transfer. While FoxP3 levels of the SMARTA TCR^pos^ T_reg_ are similar to pre-adoptive transfer donor T_reg_ after 1 day of tumor residence, these T_reg_ in MC38-GP tumors significantly lose FoxP3 expression and gain the ‘TR-T_reg_’ profile by day 7 (**Table S8, 11**). When using the top 20 differential genes in the murine TR-T_reg_ cluster from the original murine data set to generate a TR-T_reg_ score (**Table S9, 10**), we observed a progressive increase in the TR-T_reg_ score (p<2.2e-16; **Fig 3j**) together with a progressive decrease in FoxP3 expression (p<2.2e-16) over time (**Fig. 3k**). Additionally, by 7 days post-AT, TR-T_reg_ significantly upregulate the Th1-like genes *CCL5*, and *IFN*_γ_ and proliferating genes *STMN1, MKI67, and TOP2A* (**Fig. 3l**). The enhancement of proliferating genes concords with the progressive expansion in numbers of SMARTA+ T_reg_ over time in the MC38-GP tumors. In contrast, on day 1 of residence in the tumors, TR-T_reg_ express multiple resting/LN-homing genes, such as *TCF7 and SELL*, and the conventional T_reg_ genes *FoxP3* and *IL2RA* (**Fig. 3l**; **Table S11**). This type of T_reg_ differentiation has been reported in the context of autoimmune disease (*9, 47, 48*), but has not previously been demonstrated in the context of cancer.

In these analyses, it is possible that the SMARTA+ FoxP3^lo^ Th1-like population was derived from the small percentage of Foxp3^-^ T_conv_ cells present after SMARTA T_reg_ sorting (**Fig. S6d**) which could have proliferated, turned on low levels of FoxP3, and taken over the SMARTA TIL population in vivo. To rigorously address this concern, we adoptively transferred a large number (5e10^5^) of pure GP-reactive T_conv_ cells (CD4+SMARTA TCR+CD25^lo^RFP-) from CD45.1+ donor mice into CD45.2+ recipient mice bearing MC38-GP or MC38WT tumors **(Fig. S7c)**. This represented roughly 50x more tumor-reactive T_conv_ CD4+ T cells than would be contaminating the sorted TR-T_reg_ used in **Fig. 3**. Importantly, these were cells taken from the same SMARTA;RIP-GP;CD45.1;RFP-FoxP3 mice used for the T_reg_ adoptive transfers (**Fig. S6a**). In comparison to 22.8% of TR-T_reg_ expressing FoxP3 (**Fig. S6d**), virtually none of tumor-reactive CD4+ T_conv_ TIL expressed FoxP3 after 14 days in vivo (**Fig. S7d**), even though endogenous recipient derived (CD45.2, SMARTA TCR^neg^) FoxP3+ T_reg_ can be readily detected in the tumor (**Fig. S7e**). This lack of FoxP3 expression by CD4+ T_conv_ cells was also recapitulated in the spleen and tumor-draining lymph nodes (**Fig. S7f**). To further address the possibility that high-dose transgenic TCR T cell adoptive transfer could artificially promote this T_reg_ ➔ Th1-like transition (*49*), we looked at the frequency of tumor-reactive (SMARTA) TCR+ CD4+ T cells in the adoptively transferred population and in the tumor 1 day after adoptive transfer. While 92.5% of adoptively-transferred T_reg_ expressed the SMARTA (tumor-reactive) TCR, only 2.2% of the CD4+ TIL-T_reg_ were from the adoptively-transferred SMARTA TCR+ cells one day after adoptive transfer, with the remaining 97.8% being endogenous T_reg_ from the recipient mouse (**Fig. S7g**). This frequency of SMARTA TCR+ T_reg_ in the tumor after one day is consistent with prior studies of adoptively-transferred CD4+ T cells (*36*).

To further support that the cells we adoptively transferred were indeed TR-T_reg_ rather than T_conv_ cells, we performed epigenetic analysis of the T_reg_-specific demethylated region (TSDR) in the TR-T_reg_ FoxP3+ and TR-T_conv_ FoxP3^neg^ cells directly from the SMARTA RIP-GP;RFP-FoxP3 mice as well as control wild-type T_reg_ (CD4^+^CD25^hi^). The rationale behind this analysis is that T_conv_ cells will exhibit high methylation of the TSDR. Conversely, thymically-derived T_reg_ have low levels of TSDR methylation that is in accordance with the sex of the donor from which the cells were derived, because FoxP3 methylation occurs on the inactive X-chromosome (*50, 51*). Therefore, while FoxP3 methylation in male T_reg_ is typically <20%, this value in female T_reg_ ranges from 30-50% methylation. Conversely, T_conv_ cells have TSDR methylation >90% (*52*). To therefore confirm that the TR-T_reg_ obtained from our female mice were T_reg_ rather than T_conv_ cells, we analyzed the methylation status of TR-T_reg_ and TR-T_conv_ cells obtained from the same SMARTA mouse. Indeed, the TR-T_reg_ isolated from female SMARTA mice have TSDR% methylation ranging from 39.6-48.9%, consistent with known FoxP3 TSDR values in T_reg_ from female mice due to lyonization) (*53*). The TR-T_conv_ cells, however, had methylation ranging from 79.3-95.9%, consistent with known FoxP3 TSDR values in T_conv_ cells. The female donor T_reg_, which we used as a control, had methylation ranging from 55.1-69.1% (**Fig. 3m**). Together, we show that our TR-T_reg_ with a Th1-like signature have methylation patterns consistent with a typical female T_reg_ profile (*54*). We conclude that TCR-stimulation of mature self-antigen-specific T_reg_ by TAA programs these cells into a proinflammatory Th1-like immune cell phenotype.

### Tumor-reactive T_reg_ gene signature from the mouse tumor defines an orthologous subset among human NSCLC TIL-T_reg_ that is enriched in anti-PD1 responders

The robust conversion of TIL-T_reg_ from resting to a Th1-like program in MC38, which is an ICB-responsive tumor, raised the question of whether a human counterpart was associated with ICB responsiveness in our neoadjuvant cohort. To refine our search for the human ortholog to this TR-T_reg_ cluster identified in our murine tumor model, we used the gene score based on the top 20 differential genes defining the murine TR-T_reg_ cluster (**Table S9, 10**). We identified the human Th1-like/cytotoxic T_reg_ cluster (**Fig. 1a**) as the predominant population enriched for these genes (**Fig 4a, S8a**). In order to define a human transcriptional TR-T_reg_ ortholog at higher resolution, we performed refined sub-clustering on the human Th1-like/cytotoxic T_reg_ cluster and identified 6 distinct sub-clusters, one of which, SC0, was dominantly enriched in the TR-T_reg_ gene score (**Fig 4b-c, S8a, b)**. SC0 expressed high levels of *IFNG, CCL4*, and *CCL5* (**Fig. 4b,c**), which are three of the up-regulated genes associated with tumor-reactivity in murine T_reg_. SC0 also expressed multiple cytotoxic genes. It was not possible to assess antigen-specific killing activity of this population in human TIL-Treg since their cognate antigens are not defined (as in the murine system) and they express no membrane molecules that would distinguish them selectively from other Treg subsets. While *FoxP3* was expressed in all subclusters of the Th1-like/cytotoxic T_reg_ (**Fig. 4c**), *FoxP3* (p=6.8E-9) and *IL2RA* (p=1.9E-9) expression were significantly decreased in SC0 relative to all other T_reg_, while Tbet (*TBX21*) was significantly increased (p<0.001). (**Fig. 4d, Fig S8c**). Thus, SC0 has many Th1 hallmarks of the TR-Treg observed in the murine TIL-Treg. Completely opposite to the Activated (1)/OX40^hi^GITR^hi^ subset, the TR-T_reg_-score^hi^ region, SC0, was enriched in R tumors (**Fig. 4e, f**, p=0.066),. Interestingly, this enrichment in ICB responders only occurs in the tumor and not in adjacent NL.

**Figure 4.**
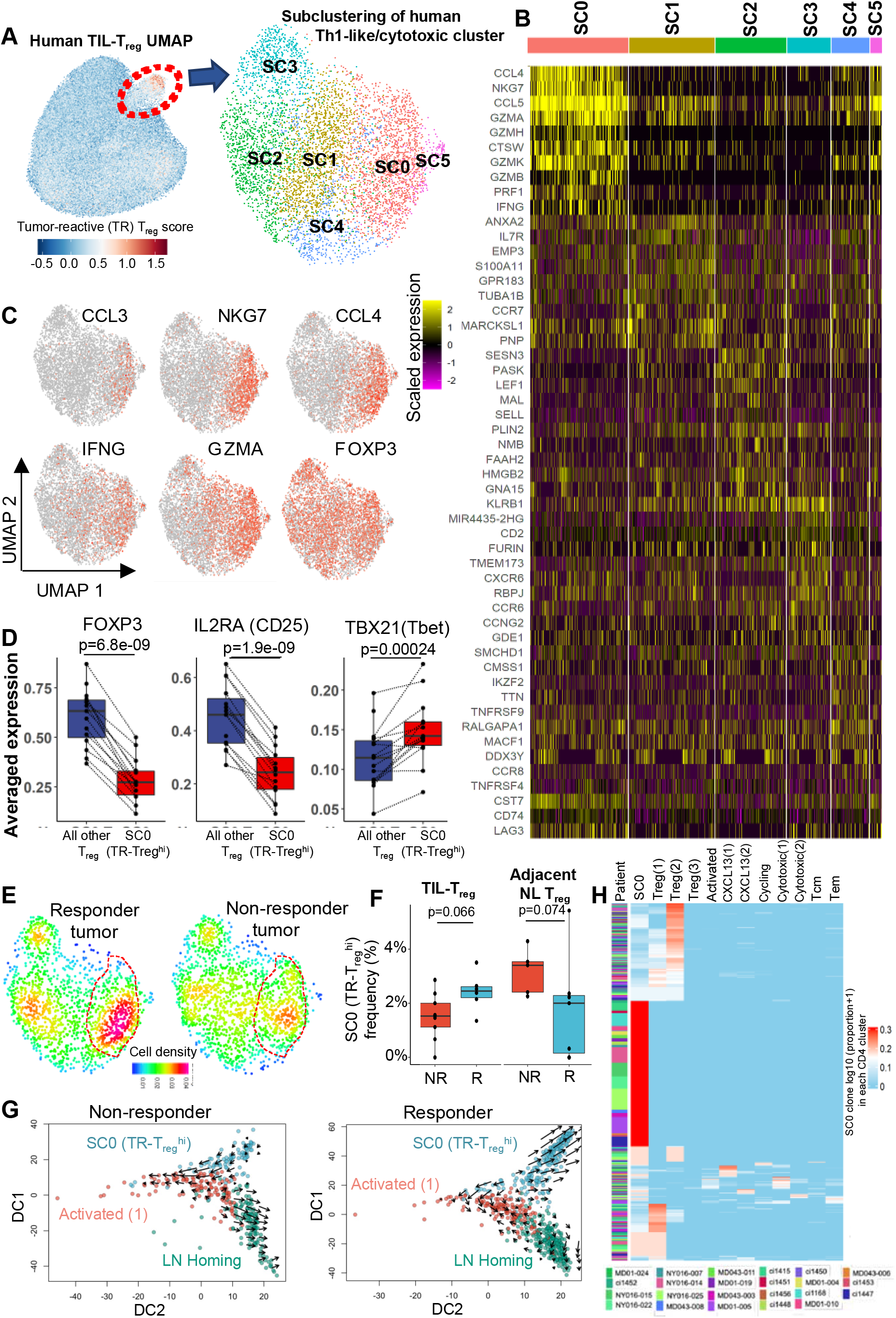
A subset of human NSCLC TIL-T_reg_ express the tumor-reactive gene signature and are enriched in anti-PD1 responding tumors. A, Red scale overlay of the tumor-reactive (TR)-T_reg_ gene score with red indicating higher expression and blue low expression. Red dotted line represents the UMAP region with highest expression. Refined clustering was performed on the Th1-like/cytotoxic subset and 2D UMAP projection of 6 unique sub-clusters (SC0-SC5) are visualized by UMAP and marked by color code. B, Relative expression of the top 10 most differential genes for each subcluster is visualized on a heatmap. C, The expression of biologically-relevant genes from the TR-T_reg_ score is visualized in red-scale. D, Boxplots showing FoxP3 (p= 6.8e-09), IL2RA (CD25; p= 1.9e-09), and TBX21(Tbet, p=0.00024) expression by TIL-T_reg_ in SC0 vs. all other T_reg_. Each dot represents an individual tumor sample/patient. Comparisons were performed at the individual patient level using paired t-test. E, Cell density plots of the Th1-like/cytotoxic T_reg_ subclusters stratified by response/treatment status. The TR-T_reg_ score^hi^ population is indicated with a dotted line. Increased density is represented by red scale and decreased density by green/blue. F, Boxplots showing the relative proportion of SC0 relative to all other T_reg_ in the tumor (left) and adjacent normal lung (right) of non-responders (red) and responders (blue). Comparisons were performed at the individual patient level using Wilcoxon rank test. G, Diffusion map with RNA velocity between SC0, Activated (1)/OX40hiGITR^hi^, and LN-homing clusters for non-responders (left) and responders (right). H, Cross-cluster sharing of T_reg_ TCR clonotypes detected in the SC0 subcluster with non-SC0 T_reg_ and T_conv_ clusters (as shown in Fig. S1c). Red indicates a higher frequency of the clone in the relevant subcluster.

To better understand the relationship between SC0 and all adjacent clusters in the global T_reg_ UMAP (Th1-like/cytotoxic, Resting, LN homing, Activated (1)/ OX40^hi^GITR^hi^, and Activated (3); **Fig. 1a**), we performed diffusion trajectory and RNA velocity. The Activated (3), Resting, and LN-homing clusters aligned along a common trajectory, with the Activated (1)/ OX40^hi^GITR^hi^ and Th1-like/cytotoxic clusters diverging along two additional trajectories (**Fig. 4g, S8c**). Because of these diffusion trajectory commonalities, and due to the relatively high clonal sharing between the Th1-like/cytotoxic and Activated (1)/OX40^hi^GITR^hi^ clusters as compared to the other UMAP-adjacent clusters (**Fig. S8d)**, we further refined this analysis to strictly assess the directionality between the LN-homing, Activated (1)/OX40^hi^GITR^hi^, and the TR-T_reg_ score^hi^ cluster (SC0). RNA velocity analysis revealed a markedly divergent direction of SC0 when comparing NR to R **(Fig. 4g)**. R SC0 T_reg_ have vectors predominantly showing net differentiation into the SC0 cluster and away from the Activated (1)/OX40^hi^GITR^hi^ and LN homing clusters **(Fig. 4g)**. We did not observe this pattern in NR. Lastly, to confirm the relationship of the Th1-like T_reg_ in the SC0 cluster to other T_reg_, rather than CD4+ T_conv_ cells, we analyzed the transcriptional heterogeneity of individual TCR clones that were found in the SC0 subcluster. Of the 441 clones that comprised SC0, 262 (59.4%) were found in at least one other CD4+ T cell cluster (Figs. 4h and S8e). Of the these ‘shared’ clones, 173 (66.0%) were solely detected in other, non-SC0 T_reg_ subsets, whereas only 43 shared clones (16.4%) were shared solely with T_conv_ subsets, and 46 clones (17.6%) were shared with both T_reg_ and T_conv_ subsets. Notably, the dominant sharing by SC0 clones with non-SC0 T_reg_ was observed across all patients, and the 16.4% of shared clones that were found in T_conv_ subsets were heterogeneous in nature, with no obvious T_conv_ subset with which most clones were shared. These analyses provide support for the relative enrichment and maintenance of the Th1-like cluster in R tumors, and the origination of these cells from T_reg_ rather than T_conv_ CD4+ TIL.

## Discussion

Using a combination of single cell transcriptional analysis of human NSCLC TIL-T_reg_ and trackable TAA-specific T_reg_ in a murine cancer model, we resolved multiple transcriptionally distinct TIL-T_reg_ subsets – 10 in human and 5 in mouse with remarkable cross-species matching. One of the activated subsets, characterized by selective expression of OX40, GITR, and high LAG3, and multiple immune suppression-associated genes such as CD39 and EBI3, was functionally suppressive *in vitro* and correlated with ICB resistance. In a murine model, T_reg_ specific for a TAA downmodulated FoxP3 and progressively developed into a population resembling Th1 cells, which was dependent on expression of cognate antigen. Using a gene signature from the murine population, we found a clear ortholog in human NSCLC TIL-T_reg_ that is more highly represented among pathologic responders to anti-PD-1.

While OX40 and GITR expression by TIL-T_reg_ have been reported for a number of human cancers (*55, 56*), our findings define a single specific subset that expresses these TNFRSF members along with the highest levels of multiple suppressive genes among the overall population. The particularly high LAG3 expression, which promotes T_reg_ suppression in murine models (*57*–*59*), on the Activated (1)/OX40^hi^GITR^hi^ subset raises the possibility that the success of LAG3 blockade in human cancer immunotherapy may be related to inhibition of suppressive function by this OX40^hi^GITR^hi^ T_reg_ population. Furthermore, it is notable that a major source of OX40L in the tumor microenvironment is the neoantigen-specific CD8 population, with much higher expression in TIL from anti-PD-1-resistant tumors. These findings, together with the strong proliferative and T_reg_ activation programs among the NSCLC TIL-T_reg_ induced by OX40L and previous evidence that OX40 signaling stabilizes T_reg_ (*24*), support the notion that tumor-specific CD8 cells can directly enhance T_reg_ suppression as a form of feedback inhibition. If so, the agonistic OX40 antibodies under clinical investigation may be counterproductive; rather, OX40 pathway blockade might be considered as an immunotherapy strategy.

The remarkable similarity among TIL-T_reg_ subsets between a commonly used murine cancer and human NSCLC was validated by comparative gene set enrichment analysis. The most unexpected finding from our murine model is that when cognate antigen is expressed in the tumor, tumor-reactive T_reg_ accumulate, down-modulate FoxP3, and progressively develop from a resting state into a population expressing Tbet, Th1-like cytokines and chemokines, as well as multiple cytotoxic molecules. We rigorously ruled out the possibility that the TAA-specific Th1-like T_reg_ population comes from expansion and intra-tumoral accumulation from the few (<5%) TAA-specific T_conv_ CD4 cells contaminating the T_reg_ at the time of adoptive transfer that might transiently up-regulate FoxP3. The development of this population in an inflamed tumor (representative of half of human NSCLC) is reminiscent of “ex-T_reg_” that develop in autoimmune settings and can exacerbate auto-inflammatory responses. Although these cells have an established functional role in the precipitation of autoimmune disease, their contribution to antitumor immunity has not previously been studied. Apropos of this notion, the instability of FoxP3 expression by T_reg_ is well-established (*60*), but its association with ‘ex-Treg’ phenotype and function has not been previously reported. In the present study, we find that virtually all the TAA-specific T_reg_ make this conversion in the MC38 tumors over a 2-week period, in contrast to the minority of T_reg_ reported to convert to ex-T_reg_ in autoimmunity settings (48-52). Th1-like Tbet^+^ T_reg_ have been shown to develop in inflamed tissues. In fact, the expression of Tbet and IFNγ by T_reg_ is required for the development of autoimmune colitis in murine models (*41*) and human multiple sclerosis(*42*), and this subset is enriched in type 1 diabetic patients (*38*). These prior findings are consistent with the higher incidence of immune-related adverse events in immunotherapy-sensitive tumors (*61*). Specifically, tumor-reactive T_reg_ may contribute positively to anti-tumor immunity while also facilitating development of therapy-induced autoimmunity.

By generating a gene profile for these tumor-specific T_reg_ in the mouse, we identified a human TIL-T_reg_ ortholog, which, in contrast to the OX40^hi^GITR^hi^ subset, is more highly represented among anti-PD-1-sensitive tumors. The presence of distinct T_reg_ subsets with putatively opposing function regarding anti-tumor immunity begs the key question of which signals in the tumor microenvironment drive incoming T_reg_ to one or the other population. Elucidation of these signals will likely define important targets for immunotherapy intervention. Another key endeavor is definition of the antigen-specificity of the human T_reg_ subsets. In clear contrast to a recent study of melanoma TIL-T_reg_ (*62*), our analysis of the more highly represented clones in the Th1-like TIL-T_reg_ subset via TCR gene transfer and screening of predicted MHC2-binding neoantigenic peptides failed to reveal any such recognition **(Fig S8f)**. Fundamentally, one might indeed expect that tumor recognition by T_reg_ would preferentially target self-antigens rather than mutation-associated neoantigens, since the natural T_reg_ repertoire is positively selected on self-antigens in the thymus (*28, 29*). Because TAA-specific T_reg_ in the murine model differentiate exclusively into a Th1-like program rather than the activated OX40^hi^GITR^hi^ program, we postulate that this suppressive subset may enter the tumor having already developed a suppressive program due to recognition of self-antigen not expressed by the tumor. The dramatically different clonality between melanoma and lung cancer concords with the apparent differences in frequency of neoantigen-specific T_reg_ observed by Oliveira et al in melanoma as opposed to our findings in lung cancer (*62*). We postulate that the distinct repertoires of T_reg_ antigen recognition in melanoma vs lung cancer reflects the different microenvironmental cues between the two tumor types, as witnessed by their different responses to immunotherapy. Moreover, as combination ICB becomes more standard, it is likely that different therapeutic regimens will differentially impact TIL-T_reg_ function.

## Acknowledgments

We thank the Experimental and Computational Genomics Core (ECGC) and the FEST and TCR Immunogenomics Core at the Sidney Kimmel Comprehensive Cancer Center. The Johns Hopkins Upper Aerodigestive Biorepository and the Genetic Resources Core Facility (GCRF). K. Maly, L. Hartman, R. Carlson and our respective administrative teams, as well as clinical support from Iiasha Beadles and Chanice Barkley. We would like to thank Dr. Matthais von Herrath at the La Jolla Institude for Allergy and Immunology for kindly providing the RIP-GP mice and Dr. Remy Bosselut at the National Cancer Institute, NIH for providing the MC38-GP tumor line.

## Author contributions

Conceptualization: AGD, JZ, LSC, DMP, KNS

Methodology: AGD, JZ, CMC, BZ, TL, JXC, LSC, MN, RH, MM, SB, DS, RR, AT, NI, DV, JS

Investigation: AGD, JZ, CMC, BZ, TL, JXC, LSC, MN, RH, MM, SB, SC, AM, CS, LT, LT

Data Analysis and Visualization: AGD, JZ, CMC, TL, BZ, HJ

Funding acquisition: KNS, DMP, HJ

Supervision: KNS, DMP, HJ, SY, PMF, JT, JRB, VA

Writing – original draft: AGD, JZ, KNS, DMP

Writing – review & editing: AGD, JZ, SY, KNS, DMP, HJ

## Data and materials availability

All processed data and code will be made readily available without restriction upon publication. Due to the sensitive nature of genomic data, raw single cell transcriptomic data will be available upon signature of a data use agreement as mandated by Johns Hopkins University. All materials will be shared with the scientific community to enable reproduction and/or validation of our results.

## Supplementary Materials

Materials and Methods

Figs. S1 to S8

Tables S1 to S13

References 63–79

