## Supplementary figures and methods for "Lung tumor-infiltrating T_reg_ have divergent transcriptional profiles and function linked to checkpoint blockade response"

**This PDF file includes:**

Materials and Methods

Figs. S1 to S13

Materials and Methods

Patients and Biospecimens

This study was approved by the Institutional Review Boards (IRB) at Johns Hopkins University (JHU) and Memorial Sloan Kettering Cancer Center and was conducted in accordance with the Declaration of Helsinki and the International Conference on Harmonization Good Clinical Practice guidelines. The patients described in this study provided written informed consent. All biospecimens were obtained from patients with stage I-IIIA NSCLC who were enrolled in a phase II clinical trial evaluated the safety and feasibility of administering two doses of anti-PD-1 (nivolumab) before surgical resection. Pathological response of primary tumors were done as previously reported (*17*, *63*). Tumors with no more than 10% residual tumor cells were considered to have an MPR.

Coupled single cell TCRseq/RNAseq

Cryobanked T cells were thawed and washed twice with pre-warmed RPMI with 20% FBS and gentamicin. Cells were resuspended in PBS and stained with a viability marker (LIVE/DEAD Fixable Near-IR; ThermoFisher) for 15 min at room temperature in the dark. Cells were the incubated with Fc block for 15 min on ice and stained with antibody against CD3 (BV421, clone SK7) for 30 min on ice. After staining, highly viable CD3^+^ T cells were sorted into 0.04% BSA in PBS using a BD FACSAria II Cell Sorter. Sorted cells were manually counted using a hemocytometer and prepared at the desired cell concentration (1,000 cells per μl), when possible. The Single Cell 5′ V(D)J and 5′ DGE kits (10X Genomics) were used to capture immune repertoire information and gene expression from the same cell in an emulsion-based protocol at the single-cell level. Cells and barcoded gel beads were partitioned into nanolitre-scale droplets using the 10X Genomics Chromium platform to partition up to 10,000 cells per sample followed by RNA capture and cell-barcoded cDNA synthesis using the manufacturer’s standard protocols. Libraries were generated and sequenced on an Illumina NovaSeq instrument using 2 × 150-bp paired end sequencing. 5′ VDJ libraries were sequenced to a depth of ~5,000 reads per cell, for a total of 5 million to 25 million reads. The 5′ DGE libraries were sequenced to a target depth of ~50,000 reads per cell.

Data preprocessing and quality control in human single-cell data

Cell Ranger v3.1.0 (*64*) was used to demultiplex the FASTQ reads, align them to the GRCh38 human transcriptome, and extract their “cell” and “UMI” barcodes. The output of this pipeline is a digital gene expression (DGE) matrix for each sample, which records the number of UMIs for each gene that are associated with each cell barcode. The quality of cells was then assessed based on (1) the number of genes detected per cell and (2) the proportion of mitochondrial gene/ribosomal gene counts. Low-quality cells were filtered if the number of detected genes was below 250 or above 3× the median absolute deviation away from the median gene number of all cells. Cells were filtered out if the proportion of mitochondrial gene counts was higher than 10% or the proportion of ribosomal genes was less than 10%. For single-cell VDJ sequencing, only cells with full-length sequences were retained. Dissociation/stress associated genes (*65*, *66*), mitochondrial genes (annotated with the prefix “MT-“), high abundance lincRNA genes, genes linked with poorly supported transcriptional models (annotated with the prefix “RP-“) and TCR (TR) genes (TRA/TRB/TRD/TRG, to avoid clonotype bias) were removed from further analysis. In addition, genes that were expressed in less than five cells were excluded.

Single cell data integration and clustering

Seurat (3.1.5) was used to normalize the raw count data, identify highly variable features, scale features, and integrate samples. Principal component analysis (PCA) was performed based on the 2,000 most variable features identified using the vst method implemented in Seurat. Immunoglobulin genes and specific mitochondrial related genes were excluded from clustering to avoid cell subsets driven by the above genes. Dimension reduction was done using the RunUMAP function. Harmony (*67*) was used to integrate across samples/batches. Cell markers were identified by using a Wilcoxon rank sum test. Genes with adjusted p.value < 0.05 were retained. Clusters were labeled based on the expression of the top differential gene in each cluster as well as canonical immune cell markers. Global clustering on all CD3 T cells and refined clustering on CD4 T cells, Treg cells and cytotoxic Treg cells were performed using same procedure. CD4+ T cells were selected by including all clusters with high CD4 expression and excluding CD8+ population (*2*). Treg cells and cytotoxic Treg cells were selected based on top differential genes from clustering.

Single cell subset pseudobulk gene expression analysis

PCA was performed on a standardized pseudobulk gene expression profile, where each feature was standardized to have a mean of zero and unit variance. In Treg PCA, we first aggregated read counts across cells within each sample to produce a pseudobulk expression profile and normalized these pseudobulk expression profiles across samples by library size. Linear regression was applied to address potential batch effects on the normalized pseudobulk profile. Batch corrected pseudobulk gene expression profile was then averaged across multiple replicates (e.g., batches) for each biological sample. Highly variable genes (HVGs) were selected by fitting a locally weighted scatterplot smoothing (LOWESS) regression of standard deviation against the mean for each gene and identifying genes with positive residuals. Samples were embedded into the PCA space based on these selected HVGs and canonical correlation (*68*, *69*) between the first two PCs (i.e., PC1 and PC2) and a covariate of interest (i.e., tissue type or response status or treated status) was calculated. Permutation test was used to assess the significance by randomly permuting the sample labels 1,000 times.

Differential expression tests among Treg cell subsets

Differential expression (DE) tests for Treg cell subsets were performed using FindAllMarkers functions in Seurat with Wilcoxon Rank Sum test. Genes with > 0.25 log2-fold changes, at least 25% expressed in tested groups, and Bonferroni-corrected p values < 0.05 were regarded as significantly differentially expressed genes (DEGs). Activated (1) Treg marker genes were identified by applying the DE tests for upregulated genes between cells of Activated (1) cluster to all Tregs in the dataset and visualized using volcano plot. Same approach as applied to SMARTA+ Tregs in day 7 vs day 1.

Analysis of gene pathway enrichment in Treg subsets

Raw count gene expression was scaled and centered using the “ScaleData” function in Seurat. A composite score for selective gene pathways was computed using the AddModuleScore function and subsequently visualized in heatmap using the pheatmap (*70*) package.

Treg activation score generation and comparisons between response/treatment groups

To characterize activated Tregs, top three differentially expressed genes in activated (1) clusters relative to all other Tregs: TNFRSF18, TNFRSF4 and LAG3, were used to compute the activated Treg score using AddModuleScore function in Seurat. Activation score positive was defined as an activated Treg score >0. The frequency of Activation score positive Tregs between responders, non-responders and treatment naïve patients were compared using Wilcoxon rank-sum test and BH method adjusting for multiple testing.

RNA velocity-based differentiation trajectory tracing

The RNA velocity analysis was performed by first recounting the spliced reads and unspliced reads based on aligned bam files of scRNA-seq data using the velocyto python package. The calculation of RNA velocity values for each gene in each cell and embedding RNA velocity vector to low-dimension space were done using the SeuratWapper workflow for estimating RNA velocity using Seurat (<https://github.com/satijalab/seurat-wrappers/blob/master/docs/velocity.md>). The first two diffusion components from Diffusion map were used to construct the coordinates along with velocity for shared clones of activated (1) and activated (3) Treg clusters. Same procedure was performed for SC0 (TR Treg^hi^) cluster, activated (1) cluster and the LN homing cluster.

TR Treg score generation and comparisons between response groups

To map SMARTA+ signature from mouse Tregs to human Tregs, top 20 differentially expressed genes (DGEs) in SMARTA+ cluster were used to compute the TR Treg score in human Tregs using AddModuleScore function in Seurat. The frequency of TR Treg score^hi^ Treg cluster (defined as cluster with the highest TR Treg signature) relative to all Tregs were compared among responders and non-responders. Same procedure was performed by taking the top 20 DGEs from each mice Treg cluster to map the mice Treg signatures to human Tregs. The mice Treg cluster scoring was averaged at each human Treg cluster level and then visualized in heatmap.

Human ex vivo T_reg_ suppression assay

Cryopreserved tumor infiltrating lymphocytes (TIL) were thawed in prewarmed 20% FBS RPMI. Cells were stained with LIVE/DEAD^TM^ Fixable Aqua Dead Cell Stain (Invitrogen, L34957), CD3 (BV605, BD, 563219), CD4 (APC, BioLegend, 300552), CD8 (BB700, BD, 566452), CD127 (BB515, BD, 564423), CD25 (PE, Biolegend, 356104), OX40 (PeCy7, Biolegend, 350012), 41BB (BV421, BD, 564091), and GITR (BV786, BD, 747661). Healthy donor PBMC were used as a source for Tconv and stained with LIVE/DEAD^TM^ Fixable Aqua Dead Cell Stain (Invitrogen, L34957), CD3 (BV605, BD, 563219), CD4 (APC, BioLegend, 300552), CD8 (BB700, BD, 566452), CD127 (BB515, BD, 564423), CD25 (PE, BioLegend, 356104), CD45RA (BV786, BD, 563870), CD45RO (BV421, BD, 562649). CD3+CD4+CD8-CD25^hi^CD127^lo^OX40^hi^GITR^hi^ and live CD3+CD4+CD8-CD25^hi^CD127^lo^OX40^lo^GITR^lo^ Tregs were sorted from TIL. Live CD3+CD4+CD8-CD25-CD45RA+CD45RO- naïve Tconv were sorted from healthy donor PBMC. For division tracking, Tconvs were labeled with CellTrace^TM^ Violet (Invitrogen) as described above. Cells were cultured in X-VIVO^TM^ 15 (Lonza) with 15% Human AB Serum and 1% L-glutamine (Gibco) as previously described (*71*). The Treg suppression assay was performed as follows: 25,000 Tconvs were co-cultured with a varying ratio of Tregs in 96-well Ubottom plate. Anti-CD3/anti-CD28 coated microbeads (Dynabeads Human T Activator, ThermoFisher, 11161D) were added at a 1:16 (bead:cell) ratio with final number of cells in each well. Proliferation was analyzed by flow cytometry done on a BD FACSCelesta^TM^ after 4 days. A minimum number of 1500 CellTrace^TM^ Violet+ Tconv cells were collected and analyzed with FlowJo version 10.

TNFRL-expressing cell lines

*Generation of lentiviral transfer plasmids*

Plasmids encoding *TNFSF4 (OX40L,* Sino Biological, HG13127-UT)*, TNFSF9 (41BBL*, Sino Biological, HG15693-UT)*,* and *TNFSF18 (GITRL*, Sino Biological, HG16080-UT) were purchased. Lentiviral transfer plasmids pLenti CMV Puro 2.0 MCS were a gift from Andrew Holland (Johns Hopkins University, MD). TNFRL cDNAs were subcloned into the pLenti CMV Puro 2.0 MCS lentiviral transfer plasmid using Gibson assembly with the addition of two N-terminal, cytoplasmic flag tags. Proper insertion of the TNFRLs into the lentiviral transfer plasmids was verified by sequencing.

*Production of TNFRL lentivirus*

To produce lentiviruses, HEK293T (ATCC, Manassas, VA) cells were thawed from liquid nitrogen and cultured in Dulbecco’s modified eagle medium (DMEM, Invitrogen) containing 10% fetal bovine serum (FBS, Sigma) supplemented with L-glutamine (Gibco), 10mM HEPES (Gibco), and 100U/ml penicillin 100ug/ml streptomycin (Quality Biologic). For lentiviral production, HEK293T cells were transfected with a TNFRL lentiviral transfer plasmid, psPAX2 (Addgene plasmid #12260) and pMD2.G (Addgene plasmid #12259) at a ratio of 1:0.75:0.25, respectively, with Lipofectamine 3000 (Invitrogen) according to manufacturer’s instructions. 24 hours post-transfection the media was replaced and supernatants containing lentivirus were harvested 48 and 72-hours post-transfection. Following harvest, viral supernatants were filtered with a .45 uM filter to remove any cellular debris and were concentrated five-fold using RetroX Concentrator (Takara Bio) according to manufactures protocol. Concentrated lentivirus was stored at -80°C.

*Generation of TNRFL expressing HEK293T cells*

HEK293T cells (1x10^5^) were plated onto poly-D-lysine coated 24-well plates. The next day lentiviruses were thawed on ice and cells were transduced with lentivirus with an MOI of 10 for each TNFRL in 1.5ml media with 8ug/ml polybrene (Millipore Corp, TR-1003-G). Plates were spun for 1.5 hours at 30°C at 2000 rpm with no brake. After the spin, cells were transferred to culture incubator. The following day, culture media was replaced. Three days after transduction, cells were validated for TNFRL expression by flow using surface TNFRL specific antibodies: OX40L (BV786, BD, 563330), 41BBL (APC, BioLegend, 311505), and GITRL (APC, Miltenyi, 130-112-974) as well as intracellular FLAG-tag staining (APC, BioLegend 637307). Transduced empty vector and TNFRL expressing cells were cultured in media with 1ug/ml puromycin for selection and passaged until >95% purity.

TNFRL-agonism of tumor infiltrating Tregs

Expression of each TNFRL or lack of expression in empty vector line was established by flow cytometry as described above on Day 0 prior to plating. 25,000 TNFRL-expressing or empty vector transduced 293T cells were seeded on Day 0 into a poly-D-lysine coated 96-well flat bottom plate. On Day 1 tumor infiltrating Tregs were sorted from cryopreserved TIL from two non-MPRs. TIL were thawed in prewarmed 20% FBS RPMI and spun at 1200 rpm for 7 mins to remove residual DMSO. Cells were stained with LIVE/DEAD^TM^ Fixable Aqua Dead Cell Stain (Invitrogen, L34957), CD3 (BV605, BD, 563219), CD4 (APC, BioLegend, 300552), CD8 (BB700, BD, 566452), CD127 (BB515, BD, 564423), and CD25 (PE, Biolegend, 356104). CD3+CD4+CD8-CD127^lo^CD25^hi^ Tregs were sorted. 20,000 Tregs were plated on TNFRL-293T cells separately or empty vector-293T cells as a negative control in X-VIVO^TM^ 15 (Lonza) with 15% Human AB Serum and 1% L-glutamine (Gibco). TNFRL-agonism was allowed to occur for 6 hours. Treg cells were then harvested and scRNA-seq/TCR-seq was carried out as described.

Mice

SMARTA-1 mice (B6.Cg-*Ptprc^a^ Pepc^b^* Tg(TcrLCMV)1Aox/PpmJ) have a CD4+ T cell repertoire that recognizes the H2-IAb restricted LCMV GP61-80 epitope as previously described (*30*). To generate mice with a sufficient number Tregs, SMARTA-1 mice were crossed to RIP-GP mice that have the LCMV GP protein expressed under the rat insulin promoter allowing for expression of these epitopes in the βcells of the pancreas as well as in the thymus for positive selection. This cross allows for thymic generation of SMARTA-1 Tregs as described (*33*, *34*). SMARTA-1;RIP-GP mice were then crossed to RFP-FoxP3 reporter mice (C57BL/6-*Foxp3^tm1Flv^*/J). These mice were used for adoptive transfer into C57BL/6J wild-type mice. SMARTA-1, RFP-FoxP3, and C57BL/6J were obtained from Jackson Laboratories (Bar Harbor, ME). RIP-GP mice were obtained from the laboratory of Dr. Matthias von Herrath (La Jolla Institute, CA). All mice were housed and bred under specific pathogen-free conditions at Johns Hopkins University Animal Care and Use Facility in Cancer Research Building I. The Institutional Animal Care and Use Facility at Johns Hopkins University approved all animal experiments.

Tumor Cell Lines

Parental MC38 (MC38WT) and MC38-GP tumor lines were grown in DMEM 10% FBS, 10mM HEPES (Gibco), L-glutamine (Gibco), 100 U/ml Penicillin, 100ug/ml streptomycin (Quality Biologic). Cells were passaged once using 0.05% trypsin before flank injection. 6x10^5^ cells were injected in 100ul PBS into each mouse flank using Intradermal TB needles (BD, 309623).

Murine ex vivo T_reg_ suppression assays

C57BL/6J mice were injected with 600,000 MC38-GP tumor cells bilaterally. On tumor growth day 14, tumors and tumor draining lymph nodes (TDLN) were harvested and pooled for Treg isolation. Spleens from CD45.1 mice were taken for isolation of naïve conventional cells (Tconv). Pooled mouse tumors were digested with tumor dissociation kit (Miltenyi, 130-096-730) and CD4 cells were positively selected (CD4 (TIL) Microbeads, Miltenyi, 130-116-475). Spleens and TDLNs were digested with digest media (RPMI (Gibco), 0.726mg/ml Collagenase I (Gibco, 17018029), 0.2mg/ml DNAse (Roche, 10104159001), FBS (Sigma)) and CD4 cells were positively selected (Miltenyi, 130-117-043). CD4+ cell suspensions were stained with LIVE/DEAD^TM^ Fixable Aqua Dead Cell Stain (Invitrogen, L34957) for 20 minutes at room temperature washed and then incubated with purified Rat Anti-Mouse CD16/CD32 (Mouse BD Fc Block^TM^) for 10 mins at 4°C followed directly with surface staining by either Treg isolation panel: CD45.1 (APC-Cy7, BioLegend, 110716), CD45.2 (BV650, BioLegend, 109836) CD4 (PE-CF94, BD, 562285), CD8 (BV786, BD, 563332), CD25 (BV421, BD, 564571), OX40 (PE, BioLegend, 119410) , GITR (PerCP/Cy5.5, BioLegend, 126316), 41BB (APC, BioLegend, 106110) or Tconv isolation panel: CD45.1 (APC-Cy7, BioLegend, 110716), CD45.2 (PerCP-Cy5.5, BioLegend, 109828) CD4 (APC, BioLegend, 100411), CD8 (BV786, BD, 563332), CD25 (BV421, BD, 564571), CD62L (PE, BioLegend, 104408), and CD44 (AF700, BioLegend, 103026) for 20 minutes at 4°C. CD45.2+CD4+CD8-CD25^hi^OX40^hi^GITR^hi^ or CD45.2+CD4+CD8-CD25^hi^OX40^lo^GITR^lo^ Tregs were sorted from tumors and TDLNs. CD45.1+CD4+CD8-CD25-CD62L^hi^CD44^lo^ naïve Tconvs were sorted for effector cells. Cell sorting occurred on a BD FACSAria^TM^. For division tracking, Tconvs were labeled with CellTrace^TM^ Violet (Invitrogen) by incubating cells for 16 min at 37 degrees at a cell density of 10^6^/ml in PBS with 0.1% bovine serum albumin (BSA). Cells were vortexed every 4 mins of staining. Reaction was quenched in ice cold RPMI 1640 medium (Invitrogen) with 10% fetal bovine serum (FBS, Sigma). As previously described (*71*), Cells were cultured in RPMI 1640 medium (Invitrogen) supplemented with non-essential amino acids (Gibco), 1mM sodium pyruvate (Sigma), 10mM HEPES (Gibco), 100 U/ml Penicillin, 100ug/ml streptomycin (Quality Biologic), 50uM 2-mercaptoethanol (Gibco), 2mM L-glutamine (Gibco) and 10% FBS (Sigma). Cells were incubated at 37 deg C in 5% CO2. The Treg suppression assay was performed as follows: 50,000 Tconvs were co-cultured with a varying ratio of Tregs in 96-well Ubottom plate. Anti-CD3/anti-CD28 coated microbeads (Dynabeads Mouse T Activator, ThermoFisher, 11452D) were added at a 1:2 (bead: cell) ratio with final number of cells in each well. Proliferation was analyzed by flow cytometry done on a BD FACSCelesta^TM^ after 4 days. Cells were washed in PBS and stained with LIVE/DEAD^TM^ Fixable Near-IR Dead cell Stain (Invitrogen, L10119) and CD45.2 (PerCP-Cy5.5, BioLegend, 109828) to differentiate Tconv from Tregs. A minimum number of 1500 CD45.1+CellTrace^TM^ Violet+ cells were collected and analyzed with FlowJo version 10.

Tumor-specific T_reg_/T_conv_ adoptive transfer and tumor-specific T_reg_ isolation

SMARTA-1;RIP-GP;CD45.1;RFP-FoxP3 were treated with IL-2/JES6-1 for 4 consecutive days as previously described (*72*, *73*). Briefly, IL-2 (1.5ug/mouse; Peprotech; 212-12-100UG) was combined with JES6-1 (6.54ug/mouse; BioXcell; BE0043), brought to volume with PBS and incubated for 15 minutes at room temperature. 100ul of cytokine/antibody mixture was administered intraperitoneally to each mouse. On day 5, spleens and lymph nodes were harvested, digested, and CD4+ T cells were negatively selected (CD4+ T cell Isolation Kit, Miltenyi, 130-104-454). SMARTA TCR transgenic Tregs and Tconv were sorted by staining with LIVE/DEAD^TM^ Fixable Aqua Dead Cell Stain (Invitrogen, L34957) for 20 minutes at room temperature washed and then incubated with purified Rat Anti-Mouse CD16/CD32 (Mouse BD Fc Block^TM^) for 10 mins at 4°C followed directly with surface staining: CD45.1 (APC-Cy7, BioLegend, 110716), CD45.2 (PerCP/Cy5.5, BioLegend, 109828) CD4 (APC, BD, 100516), CD8 (BV786, BD, 563332), CD25 (BV421, BD, 564571), TCR Vα2 (AF700, BioLegend, 127824), and TCR Vβ8.3 (PE, BioLegend, 156304). CD45.1+CD8-CD4+CD25^hi^RFP+Vα2+Vβ8.3+ Treg cells and CD45.1+CD8-CD4+CD25^-^RFP-Vα2+Vβ8.3+ Tconv were sorted into PBS 0.1% BSA on a BD FACSAria^TM^. Cells were spun, counted, washed 1x in PBS and resuspended in 2.5-3x10^6^ cells/ml. 2.5-3x10^5^ cells were adoptively transferred in 100ul PBS via tail vein injection to CD45.2 C57BL/6J wild-type mice. The next day, 6x10^5^ MC38-GP cells were inoculated into flanks bilaterally. On tumor growth day 14, mice were sacrificed, and tumors were harvested. Tumors were digested and CD4+ T cells were positively selected as described above. SMARTA-1 (CD45.1+CD4+CD8-RFP+) and endogenous (CD45.2+CD45.1-CD4+CD8-CD25^hi^RFP-) Tregs were sorted and carried forward for scRNA-seq/TCR-seq as described above.

Mouse scRNAseq analysis

The resulting counts were further filtered on total UMI count, total feature count, and percentage of mitochondrial genes. For murine data sets, cells with less than 500 UMIs, less than 350 features, or more than 2.5% of counts from mitochondrial genes were excluded. The 2000 most variable genes were selected for input to principal component analysis where the top 50 principal components were used in downstream analyses. No batch effect correction was performed for the murine data sets, dimensional reduction was performed using the uwot (*74*) implementation of uniform manifold approximation and projection(*75*). Clustering was performed using the Seurat (*76*) implementation of shared-nearest neighbor graph construction followed by Louvaine clustering. Differential expression between groups of cells was performed using Wilcoxon rank-sum test followed by Benjamini-Hochberg correction for multiple testing. The fgsea (*77*) implementation of gene set enrichment analysis was on the results using the -log(p) * sign(FC) as a ranking metric (where p is p-value and FC is fold-change). The complete set of Hallmark gene sets as well as selected gene sets relating to Treg function were included, a list of which can be found in the supplementary GSEA tables.

Mouse SMARTA time-course

SMARTA-1;RIP-GP;CD45.1;RFP-FoxP3 were sorted as described and adoptively transferred into C57BL/6J mice bearing MC38-GP tumors (tumor growth day 7). Mice were sacrificed, tumors were harvested and SMARTA+ adoptively transferred cells were sorted 1 and 7 days after adoptive transfer. SMARTA cells from the pre-adoptive transfer, 1 day, 7 day, and 14 day time points were extracted from each data set, filtered for *Cd3e* expression greater than zero, and grouped. UCell, a gene set scoring method that provides cell-level scores independent of any data-set dependent normalization, was used to generate SMARTA scores for all cells with the previously defined SMARTA gene set. The scores and the normalized *Foxp3* expression were plotted and t-test was used to test the linear regression slope with null and alternative hypothesis for this t-test: H0: β1 = 0 (the slope is equal to zero) HA: β1 ≠ 0 (the slope is not equal to zero).

T_reg_ TSDR methylation analysis

Pre-adoptive transfer SMARTA-1 Tregs, Tconv, and C57BL/6J Tregs were sorted as described above. 1x10^5^ cells were pelleted, flash frozen in liquid nitrogen and stored at -80°C. Genomic DNA extraction, bisulfite modification, and pyrosequencing of the Treg-specific-demethylated region (TSDR; CNS2) was performed by EpigenDX (Hopkinton, MA) as described (*78*).

T_conv_ flow cytometry profiling

SMARTA Tconv cells were harvested as above and adoptively transferred into C57BL/6J MC38-GP and MC38-WT tumor bearing mice on day 8 of tumor growth. 7 days later, mouse tumors, TDLN, and spleens were digested with digest media (RPMI, 0.726mg/ml Collagenase I (Gibco, 17018029), 0.2 mg/ml DNAse (Roche, 10104159001), FBS (Sigma)). Cell suspensions were stained with LIVE/DEAD^TM^ Fixable Aqua Dead Cell Stain (Invitrogen, L34957) for 20 minutes at room temperature washed and then incubated with purified Rat Anti-Mouse CD16/CD32 (Mouse BD Fc Block^TM^) for 10 mins at 4°C followed directly with surface staining by CD45.1 (APC-Cy7, BioLegend, 110716), CD45.2 (BV650, BioLegend, 109836) CD4 (APC, BD, 100516), CD8 (BV786, BD, 563332), CD25 (BV421, BD, 564571), TCR Vα2 (AF700, BioLegend, 127824), TCR Vβ8.3 (PE, BioLegend, 156304), CD44 (PerCPCy5.5, BioLegend, 103026), CD62L (BV605, BioLegend, 104408), and FoxP3 (FITC, Invitrogen, 11-5773-83). Data was collected on a BD FACSCelesta^TM^. A minimum of 2x10^5^ CD45+ cells were captured. Data was analyzed on FlowJo version 10.

Killing Assay

MC38-GP cells were plated in a T75 flask with 100ng/ml IFNγ for 48hours and live, MHC2^hi^ cells were sorted (LIVE/DEAD^TM^ Fixable Aqua Dead Cell Stain (Invitrogen, L34957), I-A/I-E (AF700, BioLegend, 107622)) and carried forward for an additional 100ng/ml IFNγ stimulation for 48 hours until ~70% of tumor cells were MHC2+. On day 0, MHC2-expressing MC38-GP cells were harvested. 5x10^3^ tumor cells were plated in each well of a 96-well flat bottom plate with 100ng/ml IFNγ and allowed to adhere overnight. Day 1, tumor infiltrating SMARTA-1 Tregs and endogenous CD45.2 Tregs were harvested from MC38-GP tumors on tumor growth day 14. Tregs were isolated and sorted as described previously. Tregs were plated in varying ratios with MC38-GP cells for 18 hours. The next day specific tumor cell killing was evaluated by analyzing % tumor death over background.

Jurkat reporter cell line

A gBlock was created with the full TCRα and TCRβ chains separately and human constant regions and was synthesized (Integrated DNA Technologies, IDT). To generate a Jurkat reporter cell in which we could transfer our TCRs of interest, the endogenous T cell receptor (TCR) α and β chains were knocked out of a specific Jurkat line that contains a luciferase reporter driven by an NFAT-response element (Promega) using the Alt-R CRISPR system (Integrated DNA Technologies, IDT). Two sequential rounds of CRISPR knockout were performed using crDNA targeting the TCRα constant region (AGAGTCTCTCAGCTGGTACA) and the TCRβ constant region (AGAAGGTGGCCGAGACCCTC). crDNA and tracrRNA (IDT) were resuspended at 100uM with Nuclear-Free Duplex Buffer. They were duplexed at a 1:1 molar ratio according to the manufacturer’s instructions. The duplexed RNA was cooled to room temperature before mixing with Cas9 Nuclease at a 1.2:1 molar ratio for 15 minutes. 40pmols of Cas9 RNP complexed with gRNA were mixed with 500,000 cells in 20ul of OptiMEM, loaded into a 0.1cm cuvette (Bio-Rad) and electroporated at 90V and 15ms using an ECM 2001 (BTX, Holliston, MA). Cells were transferred to complete growth medium and expanded for 7 days. Limiting dilution was used to acquire single cell clones and gDNA was harvested using the Quick-DNA™96 Kit (Zymo Research, Irvine, CA). The regions flanking the CRISPIR cut sites were PCR amplified (TCRα forward primer: GCCTAAGTTGGGGAGACCAC, reverse primer: GAAGCAAGGAAACAGCCTGC; TCRβ forward primer: TCGCTGTGTTTGAGCCATCAGA, reverse primer: ATGAACCACAGGTGCCCAATTC) and Sanger Sequenced. Only TCRα^-^/β^-^ clones were selected. Complete knockout was confirmed by failure to restore CD3 expression on electroporation with only a TCRα or TCRβ chain, and successful CD3 expression on electroporation with both TCR chains.

CD8 was transduced into the TCRα^-^/β^-^ Jurkat reporter cells using the MSCV retroviral expression system (Clontech). gBlocks (IDT) encoding CD8α and CD8β chains separated by a T2A self-cleaving peptide was cloned into the pMSCVpuro retroviral vector by HiFi DNA assembly (New England Biolabs). The plasmid was then co-transfected with a pVSV-G envelope vector into the GP2-293 packaging cell line per the manufacturer’s instructions. Viral supernatant was harvested 48 hours after transfection and concentrated 20-fold using Retro-X Concentrator (Clontech). For transduction, non-tissue cultured treated 48-well plates were coated with 150 µL retronectin (Clontech) in PBS at 10 ug/mL overnight at 4°C. Plates were then blocked with 10% FBS for 1hr at RT followed by washing once with PBS. After removing PBS, viral particles and 2x10^5^ of TCRα^-^/β^-^ Jurkat reporter cells were added to each well in a total volume of 500 µL cell culture media. Plates were spun at 2000 g for 1hr at 20°C then incubated at 37°C. Selection with 1 µg/mL puromycin (Thermo Fisher Scientific) began three days later. Single cell clones were established by limiting dilution and clones were subsequently screened for CD8 expression by flow cytometry. To generate a Jurkat reporter line that expresses both CD4 and CD8, CD4 viral particles were produced and transduced into the CD8-expressing Jurkat reporter cells using similar procedures.

Jurkat TCR transfer

TCRs of interest were introduced into the CD4/CD8 TCRα^-^/β^-^ Jurkat reporter line by cloning the TCRα and TCRβ chains separately into the pCI vector (Promega) by HiFi DNA assembly (New England Biolabs). The two plasmids were co-electroporated into the TCRα^-^/β^-^ Jurkat reporter line using 4mm cuvettes (Bio-Rad) and 275V for 10ms for 3 pulses at 0.1 interval between pulses. Cells were rested in RPMI 10% FBS at 37 ° for 24 hours. TCR expression efficiency was assessed by CD3 expression using flow cytometry. After 24 hour rest, live Jurkat cells were counted and plated at 1:1 ratio with a patient matched lymphoblastoid cell line (LCL) and peptide pools (JPT and Sigma). Neoantigen peptides were plated and pooled (3-5 peptides per pool) at 50ug/ml per peptide to assess TCR reactivity to neoantigens. 20ug/ml of a peptide pool spanning the S protein of SARS-CoV-2 (JPT) was used to stimulate a known SARS-CoV-2-reactive TCR as positive control (*79*). Cells and peptide were co-cultured for 24 hours. TCR activity was assessed by NFAT-luciferase reporter readout using Bio-Glo™ Luciferase Assay System (Promega).

Statistical Software

All statistics were performed in R (version 4.2.1) and Graph Pad Prism (version 9.3.0).

17.	P. M. Forde, J. E. Chaft, K. N. Smith, V. Anagnostou, T. R. Cottrell, M. D. Hellmann, M. Zahurak, S. C. Yang, D. R. Jones, S. Broderick, R. J. Battafarano, M. J. Velez, N. Rekhtman, Z. Olah, J. Naidoo, K. A. Marrone, F. Verde, H. Guo, J. Zhang, J. X. Caushi, H. Y. Chan, J.-W. Sidhom, R. B. Scharpf, J. White, E. Gabrielson, H. Wang, G. L. Rosner, V. Rusch, J. D. Wolchok, T. Merghoub, J. M. Taube, V. E. Velculescu, S. L. Topalian, J. R. Brahmer, D. M. Pardoll, Neoadjuvant PD-1 Blockade in Resectable Lung Cancer. *N. Engl. J. Med.* **378**, 1976–1986 (2018).

63.	T. R. Cottrell, E. D. Thompson, P. M. Forde, J. E. Stein, A. S. Duffield, V. Anagnostou, N. Rekhtman, R. A. Anders, J. D. Cuda, P. B. Illei, E. Gabrielson, F. B. Askin, N. Niknafs, K. N. Smith, M. J. Velez, J. L. Sauter, J. M. Isbell, D. R. Jones, R. J. Battafarano, S. C. Yang, L. Danilova, J. D. Wolchok, S. L. Topalian, V. E. Velculescu, D. M. Pardoll, J. R. Brahmer, M. D. Hellmann, J. E. Chaft, A. Cimino-Mathews, J. M. Taube, Pathologic features of response to neoadjuvant anti-PD-1 in resected non-small-cell lung carcinoma: a proposal for quantitative immune-related pathologic response criteria (irPRC). *Ann. Oncol. Off. J. Eur. Soc. Med. Oncol.* **29**, 1853–1860 (2018).

64.	T. Stuart, A. Butler, P. Hoffman, C. Hafemeister, E. Papalexi, W. M. 3rd Mauck, Y. Hao, M. Stoeckius, P. Smibert, R. Satija, Comprehensive Integration of Single-Cell Data. *Cell*. **177**, 1888-1902.e21 (2019).

65.	C. H. O’Flanagan, K. R. Campbell, A. W. Zhang, F. Kabeer, J. L. P. Lim, J. Biele, P. Eirew, D. Lai, A. McPherson, E. Kong, C. Bates, K. Borkowski, M. Wiens, B. Hewitson, J. Hopkins, J. Pham, N. Ceglia, R. Moore, A. J. Mungall, J. N. McAlpine, S. P. Shah, S. Aparicio, T. C. I. G. C. Team, Dissociation of solid tumor tissues with cold active protease for single-cell RNA-seq minimizes conserved collagenase-associated stress responses. *Genome Biol.* **20**, 210 (2019).

66.	S. C. van den Brink, F. Sage, Á. Vértesy, B. Spanjaard, J. Peterson-Maduro, C. S. Baron, C. Robin, A. van Oudenaarden, Single-cell sequencing reveals dissociation-induced gene expression in tissue subpopulations. *Nat. Methods*. **14** (2017), pp. 935–936.

67.	I. Korsunsky, N. Millard, J. Fan, K. Slowikowski, F. Zhang, K. Wei, Y. Baglaenko, M. Brenner, P.-R. Loh, S. Raychaudhuri, Fast, sensitive and accurate integration of single-cell data with Harmony. *Nat. Methods*. **16**, 1289–1296 (2019).

2.	J. X. Caushi, J. Zhang, Z. Ji, A. Vaghasia, B. Zhang, E. H.-C. Hsiue, B. J. Mog, W. Hou, S. Justesen, R. Blosser, A. Tam, V. Anagnostou, T. R. Cottrell, H. Guo, H. Y. Chan, D. Singh, S. Thapa, A. G. Dykema, P. Burman, B. Choudhury, L. Aparicio, L. S. Cheung, M. Lanis, Z. Belcaid, M. El Asmar, P. B. Illei, R. Wang, J. Meyers, K. Schuebel, A. Gupta, A. Skaist, S. Wheelan, J. Naidoo, K. A. Marrone, M. Brock, J. Ha, E. L. Bush, B. J. Park, M. Bott, D. R. Jones, J. E. Reuss, V. E. Velculescu, J. E. Chaft, K. W. Kinzler, S. Zhou, B. Vogelstein, J. M. Taube, M. D. Hellmann, J. R. Brahmer, T. Merghoub, P. M. Forde, S. Yegnasubramanian, H. Ji, D. M. Pardoll, K. N. Smith, Transcriptional programs of neoantigen-specific TIL in anti-PD-1-treated lung cancers. *Nature*. **596**, 126–132 (2021).

68.	H. HOTELLING, RELATIONS BETWEEN TWO SETS OF VARIATES*. *Biometrika*. **28**, 321–377 (1936).

69.	W. Härdle, L. Simar, Applied multivariate statistical analysis (2003).

74.	J. Melville, uwot: The uniform manifold approximation and projection (UMAP) method for dimensionality reduction. *R Packag. version 15* (2020).

Figures S1-S8

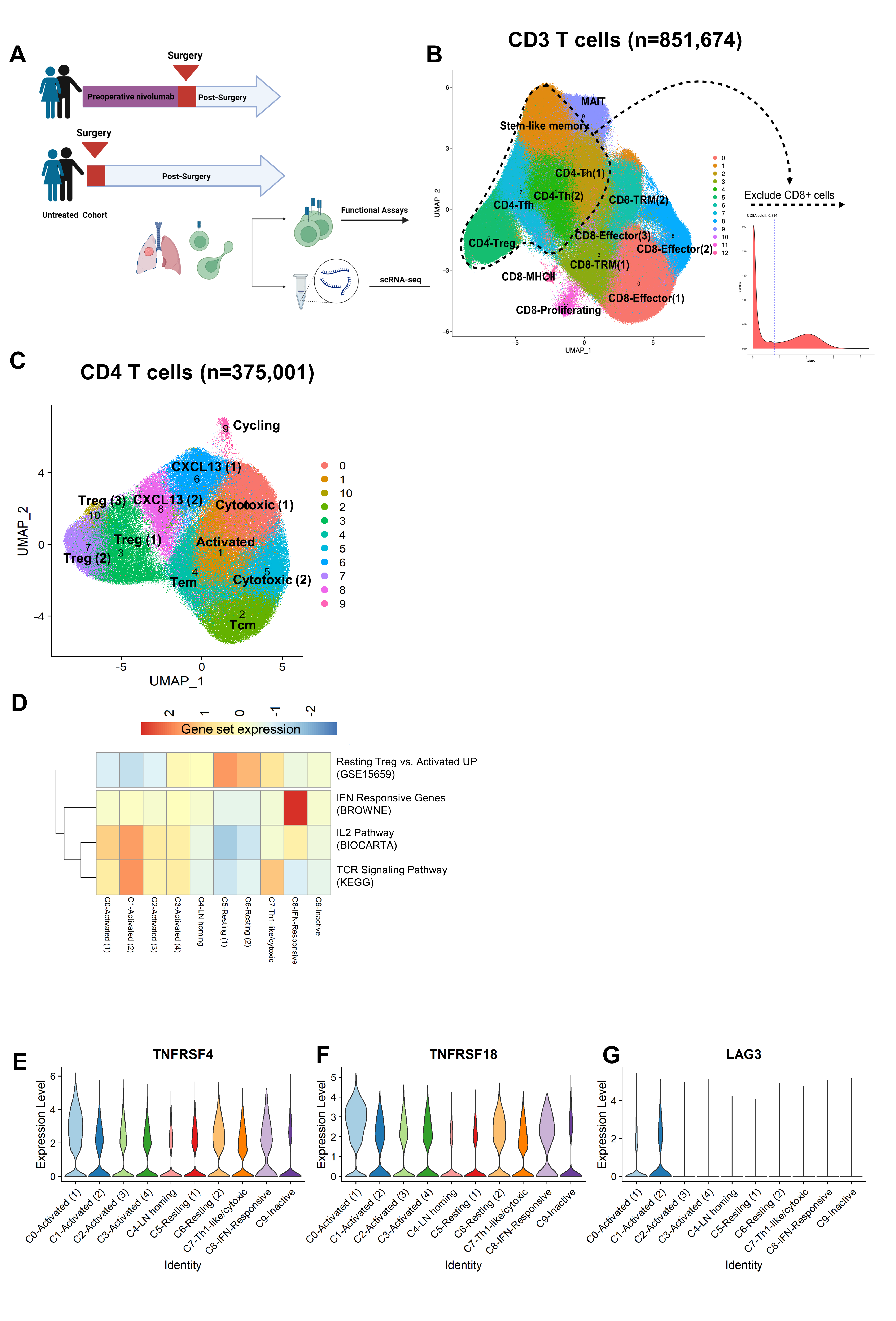

Figure S1. Study design and T_reg_ cluster definition from bulk CD3+ T cells and CD4+ T cells. Coupled single-cell (sc) RNA-seq/TCR-seq was performed on T cells isolated from resected tumor (n=15), adjacent normal lung (NL; n=12), tumor draining lymph node (TDLN; n=3), and a resected brain metastasis (n=1) from NSCLC patients treated with two doses of neoadjuvant anti-PD-1 as well as resected tumor (n=10), paired adjacent normal lung (n=8) from treatment naïve NSCLC patients. A, Schematic of study design. B, 2D UMAP projection of the expression profiles of the 851,674 T cells that passed QC. T cell subsets, defined by 13 unique clusters, are annotated, and marked by color code. C, 2D UMAP projection of the expression profiles of the 375,001 CD8 negative, CD4 positive T cells that passed QC. CD4 T cell subsets, defined by 11 unique clusters, are annotated, and marked by color code. D, Gene set enrichment revealed Activated clusters (C0-C3), Resting clusters (C5, C6), and IFN-responsive (C8). E-G Violin plots showing TNFRSF4 (OX40; E), TNFRSF18 (GITR, F), and LAG3 (G) expression by each T_reg_ sub-cluster. Comparisons were performed at the individual cell level. Colors match UMAP color code in Figure 1A.

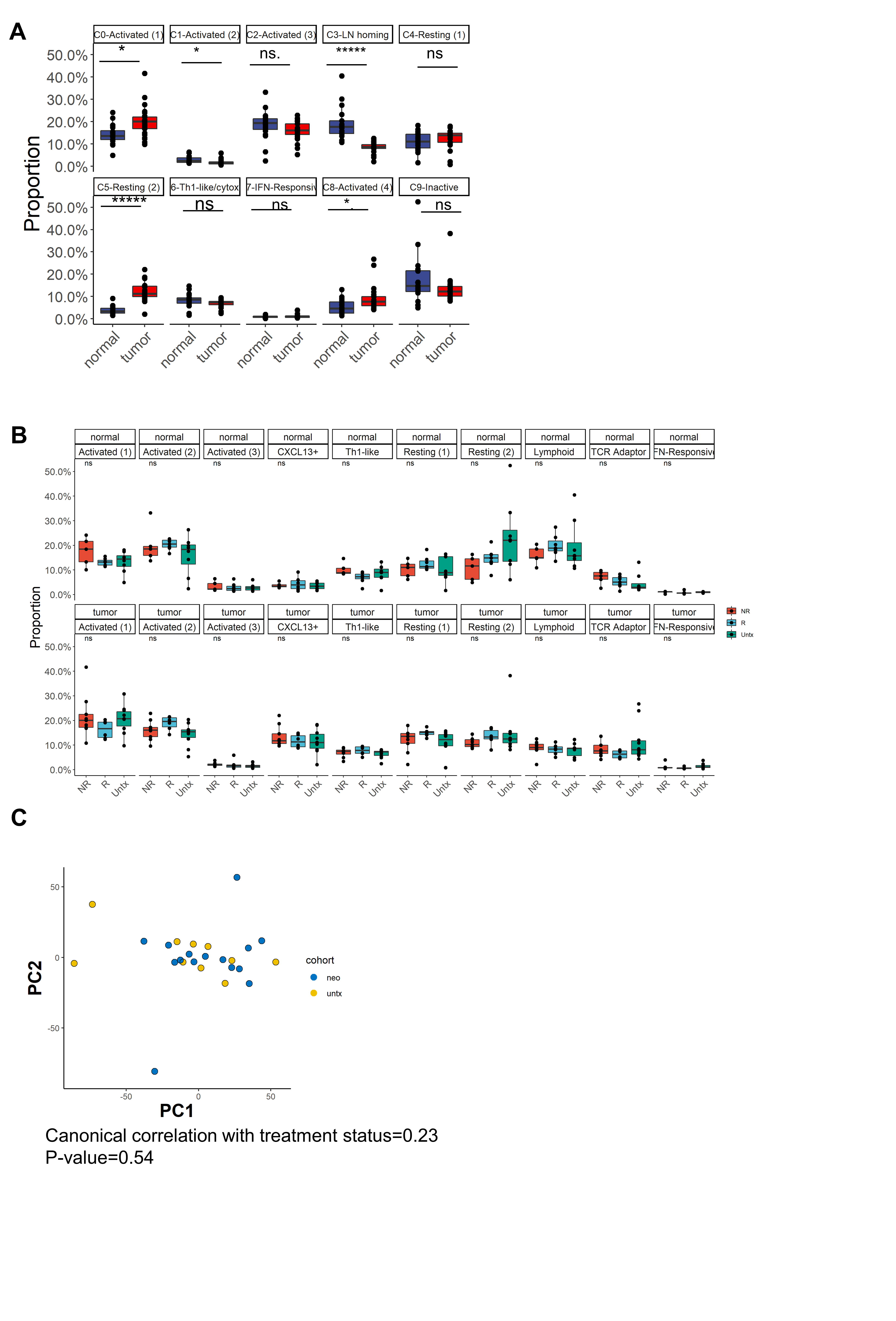

**Figure S2. T_reg_ subset proportion analysis between tumor, adjacent NL, and between treatment/response groups.** A, Differences in T_reg_ subset proportions in tumor and adjacent normal lung. *P < 0.05, **P < 0.005, ***P < 0.0005, ****P < 0.00005, *****P < 0.000005 using Wilcoxon signed-rank test, adjusted by BH method; ns, nonsignificant values. B, Differences in Treg subset proportions in tumor (bottom) and adjacent normal lung (top) stratified by treatment/response group: NR (red), R (blue), untreated (green). *P < 0.05, **P < 0.005, ***P < 0.0005, ****P < 0.00005, *****P < 0.000005 using Wilcoxon signed-rank test, adjusted by BH method; ns, nonsignificant values. C, Principal component analysis (PCA) of pseudobulk gene expression for individual samples for tumor-infiltrating anti-PD1 treated (blue, n=15; neo) and treatment-naive (yellow, n=10; untx).

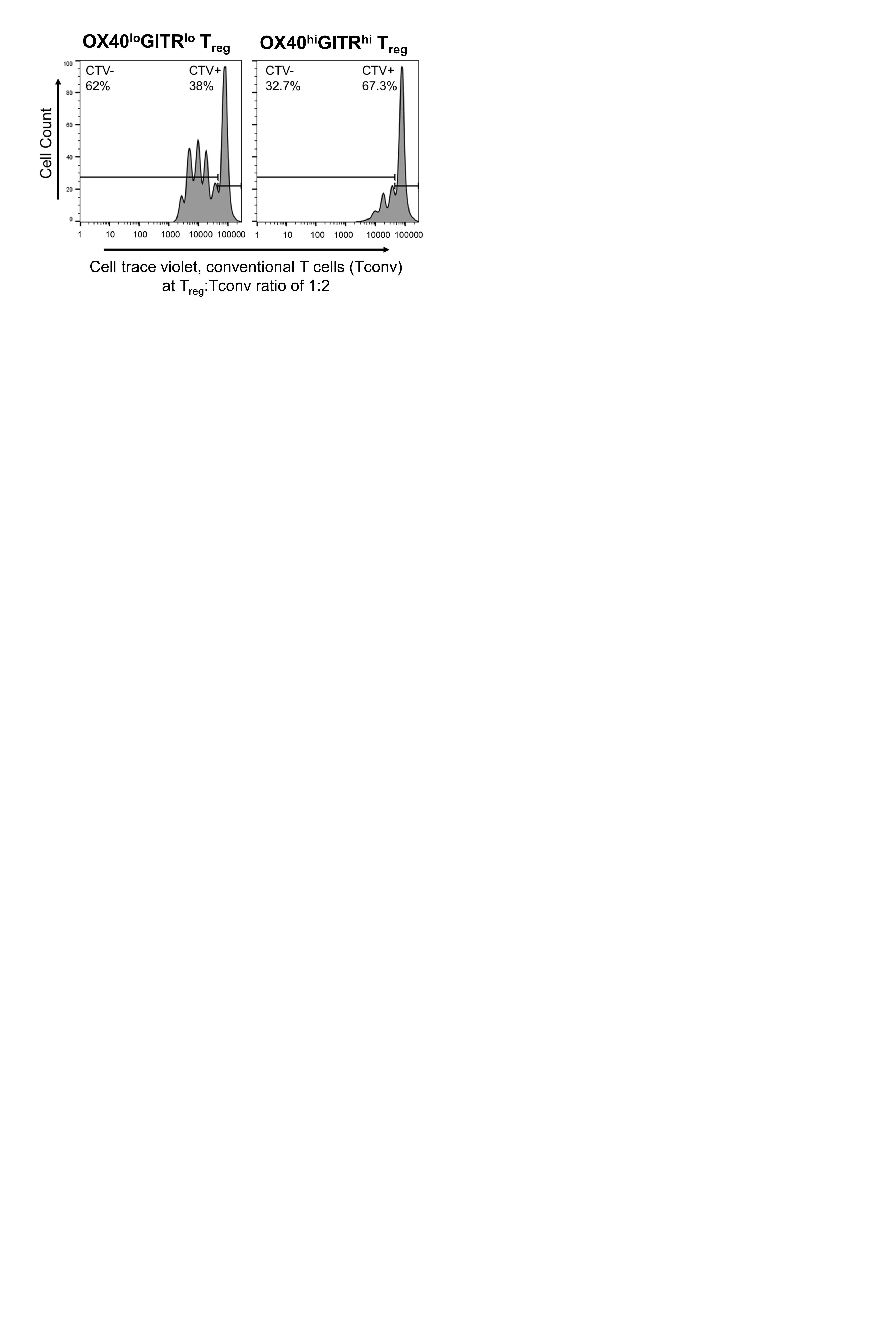

Figure S3. OX40^hi^GITR^hi^ T_reg_ suppression assays. Ex vivo suppression of CD4+ T_conv_ cells was assessed for OX40^lo^GITR^lo^ and OX40^hi^GITR^hi^ TIL-T_reg_ at 1:2 T_reg_:T_conv_ ratio as measured by Cell Trace Violet (CTV) dilution.

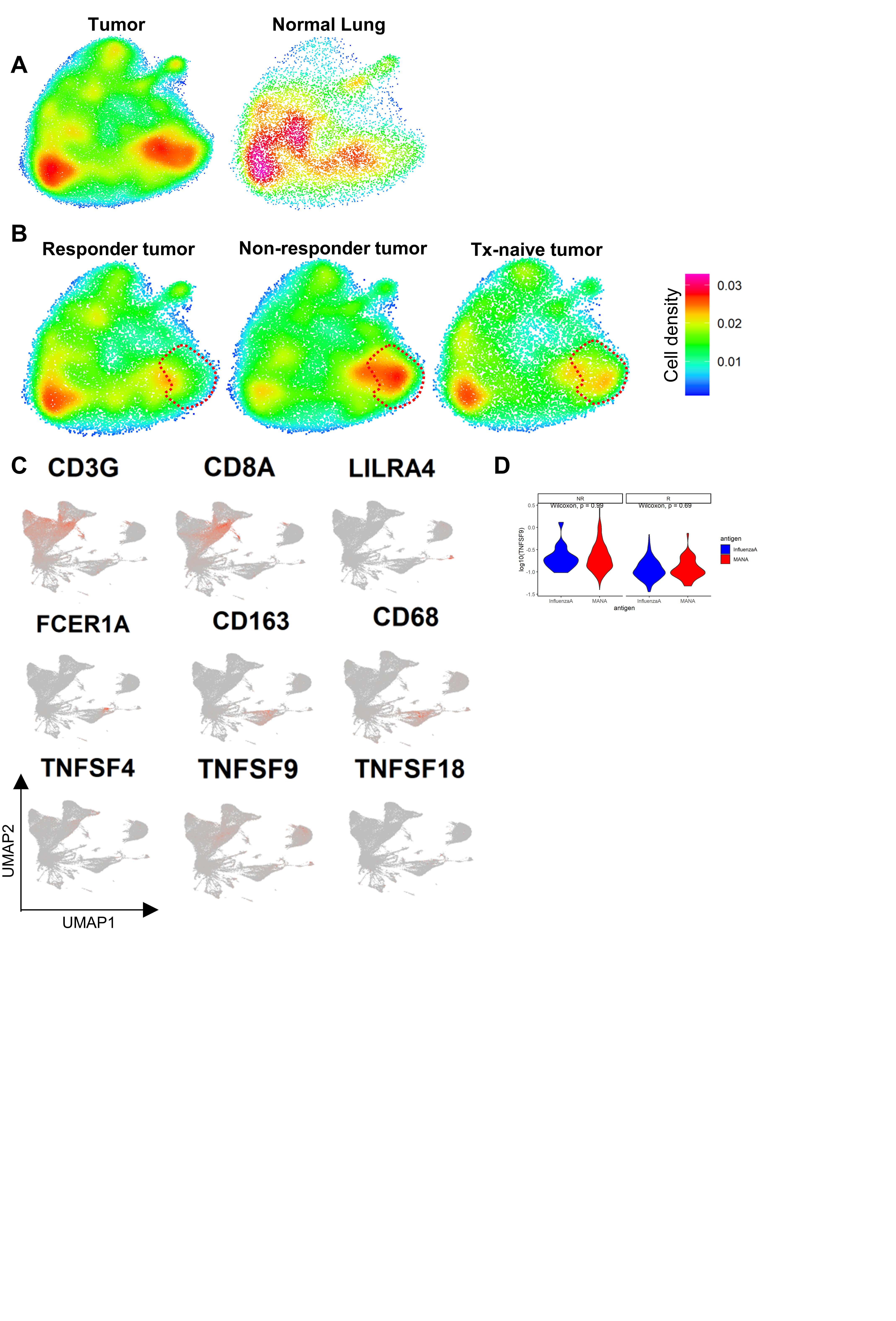

**Figure S4. Activated T_reg_ score comparisons with expression of TNFRLs on myeloid and CD8 subsets.** A, Cell density plots of T_reg_ stratified by tissue (tumor and normal lung). Increased density is represented by red scale and decreased density by green/blue. B, Cell density plots of T_reg_ stratified by response/treatment (responder, non-responder, Tx-naive). Increased density is represented by red scale and decreased density by green/blue. Red-outline corresponds to the Activated (1) OX40^hi^GITR^hi^ cluster. C, Expression of T cell and myeloid subset-defining genes and TNFSF4, TNFSF9, and TNFSF18 (OX40L, 41BBL, and GITRL, respectively) on lung TIL. D, A violin plot shows TNFSF9 (41BBL) expression by Influenza- (blue) and neoantigen (red)-specific CD8 TIL separated by NR (left) and R (right). Comparisons were performed at the individual cell level using two-sided Wilcoxon rank-sum test.

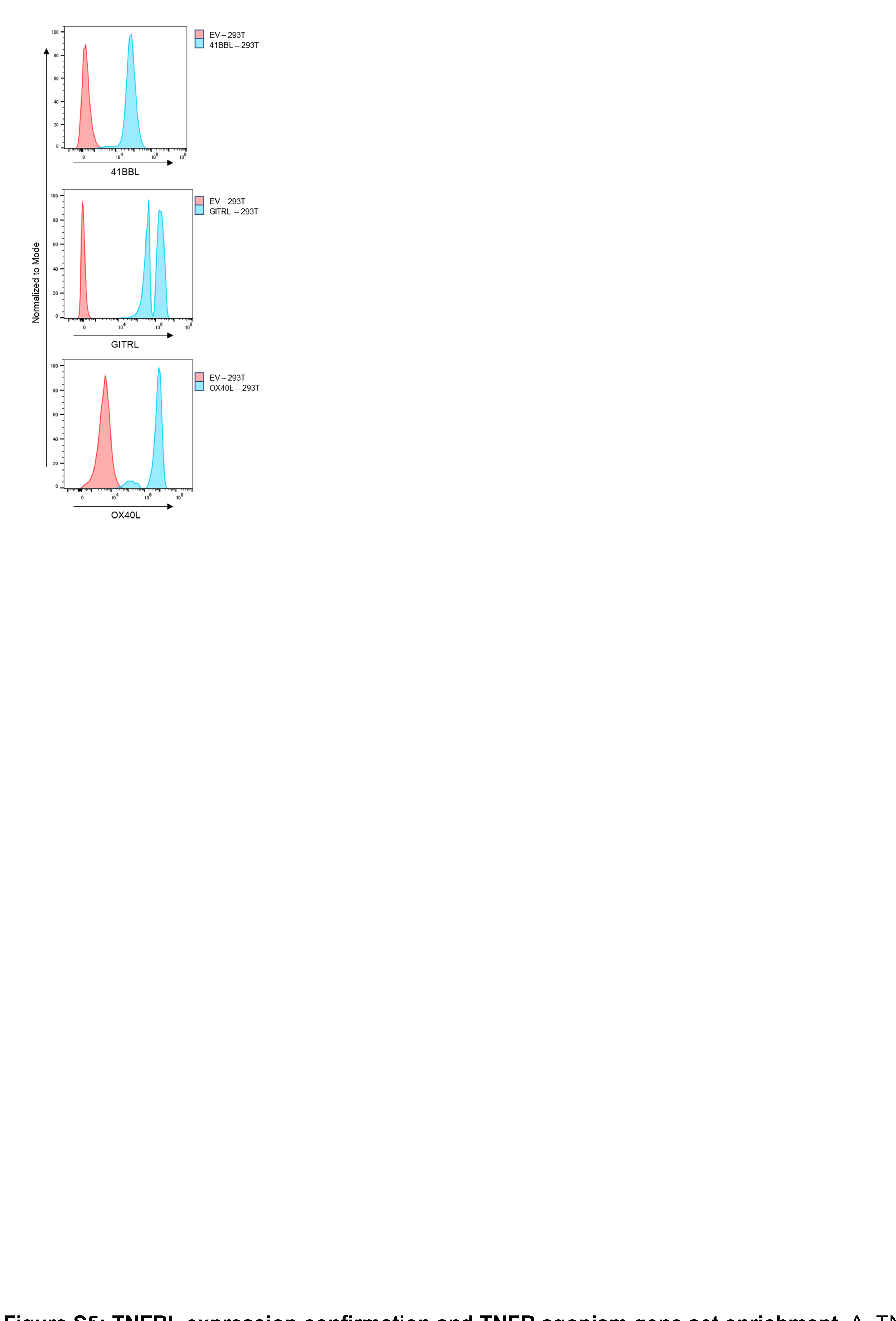

Fig S5. TNFRL expression confirmation and TNFR agonism gene set enrichment. A, TNFRL expression in HEK-293-T cell lines as compared to empty vector (EV) control. Each ligand was stained for and normalized to isotype control: (41BBL, APC; top), (GITRL, APC, middle), (OX40L, BV786, bottom).

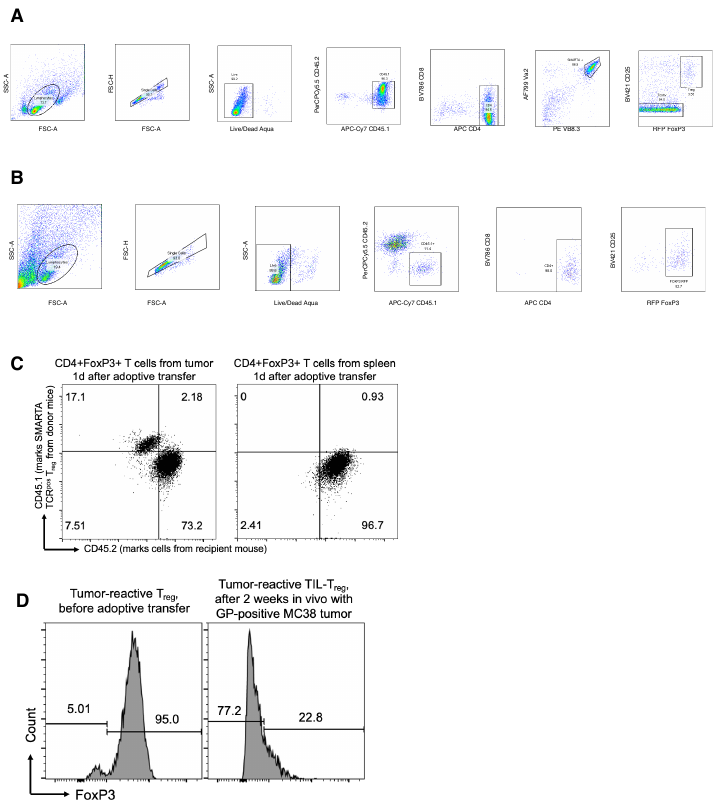

Figure S6. SMARTA T_reg_ pre and post adoptive transfer sorting and genes associated with FoxP3 differential expression. A, Sorting schematic and gating strategy showing selective sorting of CD45.1^+^CD25^hi^RFP^+^SMARTA TCR^+^ CD4^+^ T cells (SMARTA T_reg_) and CD45.1^+^CD25^lo^RFP^neg^ SMARTA TCR^+^ CD4^+^ T cells (SMARTA T_conv_) before adoptive transfer. A representative sample from 3 biological replicates is shown. B, Sorting schematic and gating strategy showing selective sorting of SMARTA T_reg_ post tumor infiltration. C, Representative example of CD45.1 and CD45.2 expression by FoxP3^+^CD4^+^ T cells from the tumor (left) and spleen (right) after 14 days in vivo with the GP-expressing MC38 tumor. D, Representative example of FoxP3 expression by tumor-reactive T_reg_ prior to adoptive transfer and after two weeks in vivo with the GP-positive MC38 tumor.

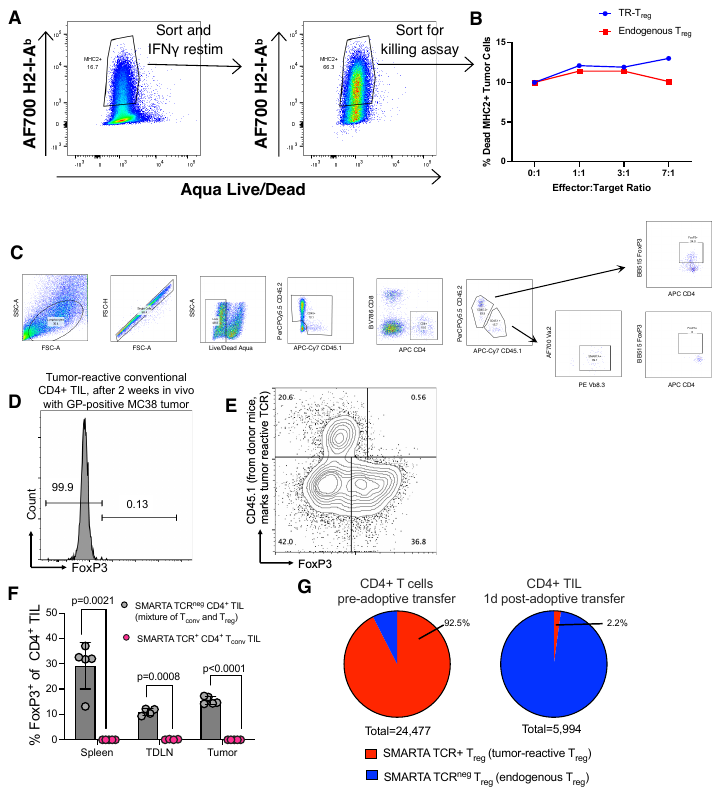

**Figure S7. SMARTA T conventional cell tumor-infiltration and flow cytometric profiling.** A, H2-I-A^b^ (MHC2)+ MC38GP tumor cells were sorted after 100ng/ml IFNγ treatment. These tumor cells were again stimulated with 100ng/ml IFNγ prior to sorting for MHC2hi tumor cells used for the in vitro killing assay. B, MHC2hi MC38GP tumor cells were co-cultured with TR-T_reg_ and endogenous T_reg_ at titrating effector to target ratios. Killing was measured by the % tumor cell death. C, Flow cytometry gating strategy to identify adoptively transferred SMARTA Tconv cells and profile them for activation and FoxP3 expression in tumor. Data are from a representative mouse (n=5). D, Comparison of tumor-reactive CD4+ TIL and TIL-Treg (RFP+) frequency from donor mice, defined by expression of CD45.1, between MC38WT (left) and MC38GP (right) after 2 weeks in vivo. E, FoxP3 expression by donor-derived (CD45.1+) and endogenous (CD45.1^neg^) tumor-reactive CD4+ T_conv_ TIL. F, Frequency of CD4+ TIL that express FoxP3 is shown for endogenous CD4+ TIL (gray) and tumor-reactive CD4+ T_conv_ TIL (pink) in spleen, tumor-draining lymph node, and tumor. Mean with SD bars are shown for each group (n=3-5). G, The frequency of CD4+ T cells expressing the SMARTA (tumor-reactive) TCR was assessed prior to and at 1 day post adoptive transfer. Data are shown as the percent of all CD4+ T cells analyzed by single cell TCRseq/RNAseq.

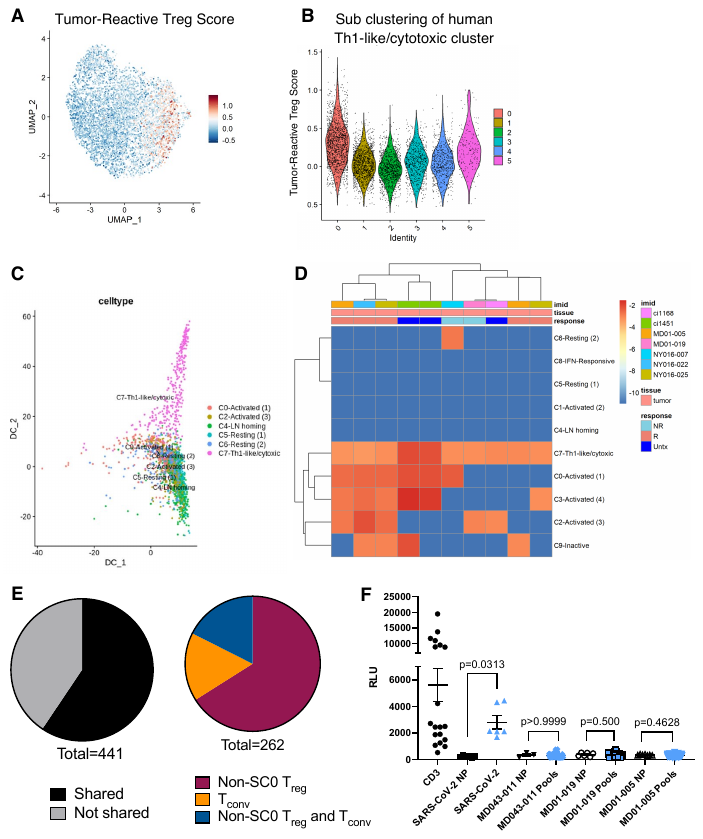

**Figure S8. TR-T_reg_ score on human T_reg_.** A, Overlay of the TR-T_reg_ score on Th1-like/Cytotoxic subclusters SC0-SC5. Red indicates higher expression; blue indicates low expression. B, Violin plots showing TR-T_reg_ score expression by subclusters SC0-SC5. Comparisons were performed at the individual cell level. C, Trajectory analysis of T_reg_ clusters selected based on the potential for trajectory relationships. D, Heatmap showing clonal sharing of Th1-like/cytotoxic T cell clonotypes with other T_reg_ subsets. Red indicates a higher frequency of the shared cloned within the relevant cluster. E, Cloning and screening of Th1-like/cytotoxic Treg against pools of putative MHC class II-restricted neoantigens. Data are shown as relative luminescence units (RLU) for selected Treg TCRs, as well as a known SARS-CoV-2-reactive TCR. All clones were also stimulated with anti-CD3 as a positive control. *P values determined using Wilcoxon signed-rank test. NP, no peptide.
